# Loss of a doublecortin (DCX) domain containing protein causes structural defects in a tubulin-based organelle of *Toxoplasma gondii* and impairs host cell invasion

**DOI:** 10.1101/069377

**Authors:** Eiji Nagayasu, Yu-chen Hwang, Jun Liu, John M. Murray, Ke Hu

## Abstract

The ~6000 species in phylum Apicomplexa are single-celled obligate intracellular parasites. Their defining characteristic is the “apical complex”, membranous and cytoskeletal elements at the apical end of the cell that participate in host-cell invasion. The apical complex of *Toxoplasma gondii* and some other apicomplexans includes a cone-shaped assembly, the “conoid”, which (in *T. gondii*) comprises 14 spirally arranged fibers that are non-tubular polymers of tubulin. The tubulin dimers used for the conoid fibers make canonical microtubules elsewhere in the same cell, suggesting that their special arrangement in the conoid fibers is dictated by non-tubulin proteins. One candidate for this role is TgDCX, which has a doublecortin (DCX) domain and a TPPP/P25-alpha domain, known modulators of tubulin polymer structure. Loss of TgDCX radically disrupts the structure of the conoid, severely impairs host cell invasion, and slows growth. The defects of TgDCX-null parasites are corrected by re-introduction of a TgDCX coding sequence.

## Introduction

Infection with *Toxoplasma gondii* can cause severe illness when the infection is contracted congenitally or when it is reactivated in immunosuppressed hosts. *T. gondii* is one of ~6,000 species of intracellular protozoan parasites in the phylum Apicomplexa, all members of which are parasitic, including various important human / animal pathogens such as *Plasmodium* (causative agents of malaria), *Cryptosporidium* (cryptosporidiosis), *Theileria* and *Babesia* (important pathogens of cattle) and *Eimeria* (pathogens of poultry and cattle),.

The apicomplexan parasites belong to the super-phylum Alveolate, organisms that have “alveola”, adjoining membrane sacs constituting two additional layers of membrane underlying the plasma membrane. Apicomplexa is one of the three major clades of Alveolate, along with dinoflagellates that diverged from the apicomplexans several hundred million years ago, and ciliates that diverged from other alveolates up to 1 billion years ago (Leander and Keeling, 2003). A number of organisms belonging to sister clades of apicomplexans and dinoflagellates have been identified in recent years. This includes marine photosynthetic relatives of the apicomplexans (chromerids), free-living relatives of the apicomplexans that prey on other cells by myzocytosis (colpodellids), and marine parasitic relatives of dinoflagellates that can be cultured in a free-living form (perkinsids). Morphologically, the perkinsids and the chromerids are unified with the apicomplexans not only by the cortical membrane sacs shared by all the alveolates, but also by a striking membrane-cytoskeletal assembly called the apical complex, which consists of electron dense, elongated vesicles associated with a cone-shaped or flattened array of tubulin polymers located at the apical end of the cell. In perkinsids and the chromerids, the apical complex contains a half-closed cone structure (“pseudoconoid”) formed of a sheet of microtubules (30–35 in the case of *Chromera velia* (Portman et al., 2014)). In *T. gondii*, the cytoskeletal apical complex includes three rings (the polar rings), a pair of short straight microtubules (the intra-conoid microtubules) and the cone-shaped conoid, a left-handed spiral of 14 fibers made from a non-tubular tubulin polymer (Figure 1). Unlike typical microtubules, conoid fibers are not closed tubes and contain only ~9 protofilaments (Hu et al., 2002).

**Figure 1.**
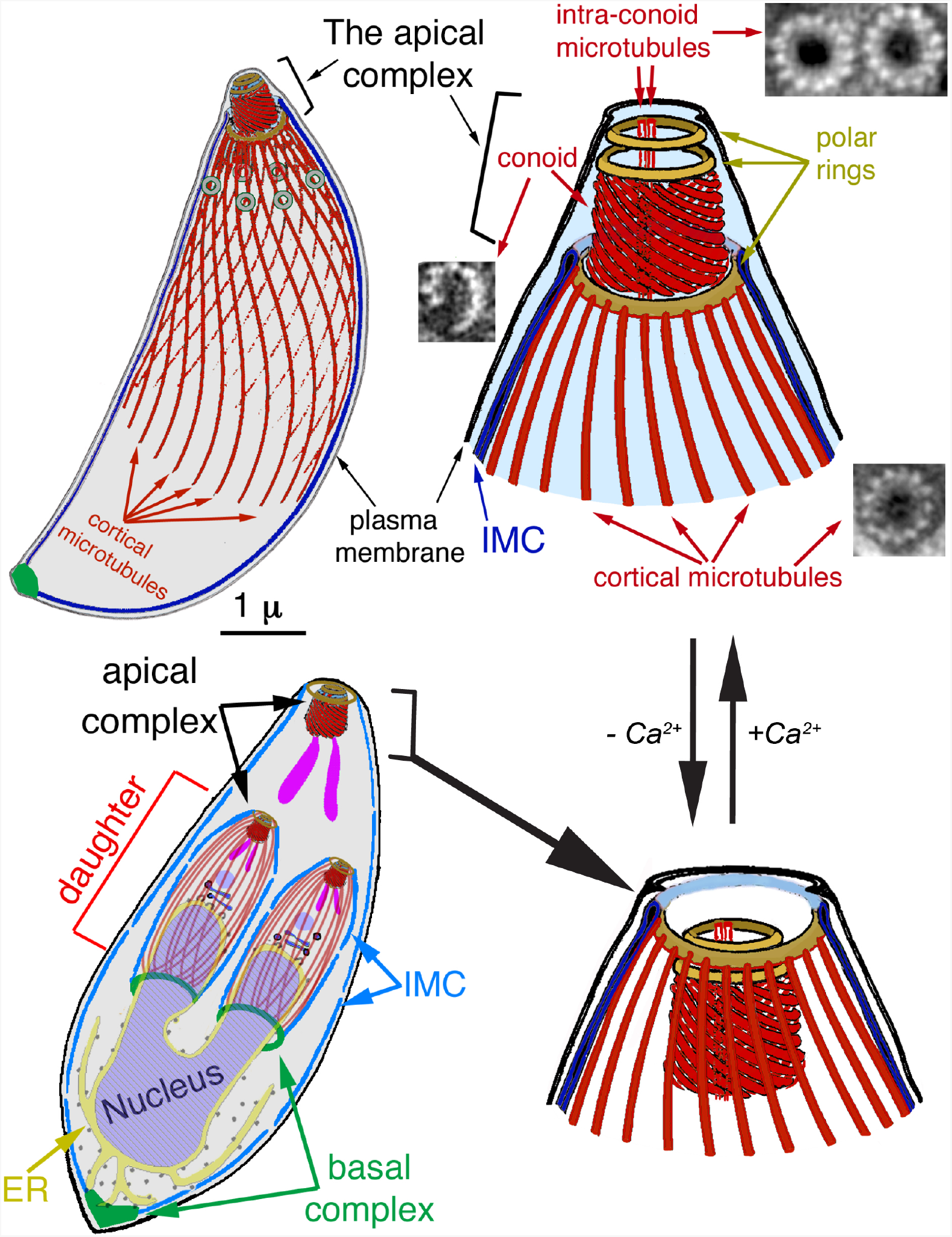
Diagram of the T. gondii cytoskeleton (from (Liu et al., 2016). The 22 cortical microtubules, 2 intra-conoid microtubules, and 14 conoid fibers, which are non-tubular tubulin polymers, are shown in red. EM images of each of those polymers are also shown in cross-section (Hu et al., 2002b). IMC: Inner Membrane Complex. Bottom left: a replicating parasite, with daughter parasites being built inside the mother. The cortical microtubules of the adult are omitted for clarity. At this stage of daughter formation, most of the membrane-bound organelles have been produced or duplicated and partitioned into daughters, except for the mitochondrion (Nishi et al., 2008), though only the Golgi stack (dark-blue) and apicoplast (a plastid-like organelle, light-purple), and rhoptries (one of the specialized organelles for invasion, purple), are shown here. In intracellular parasites, the conoid is normally retracted, as shown in the lower diagrams. An increase in cytoplasmic [Ca^2+^], which normally accompanies egress from the host cell and re-invasion, triggers extension of the conoid, as in the upper diagrams.

The tubulin dimers that form the conoid fibers are used elsewhere in the same cell to make canonical microtubules, suggesting that their special arrangement in the conoid fibers is dictated by non-tubulin proteins. Previously we partially purified the apical complex of *T. gondii* and identified its protein components (Hu et al., 2006). One of the proteins highly enriched in the apical complex fraction is TgDCX, which contains domains belonging to two protein families (“P25 alpha” and “DCX”) that are ubiquitous among metazoa. All sequenced apicomplexan genomes (Nagayasu et al., 2006; Orosz, 2009), as well as the genomes of chromerids and perkinsus (Orosz, 2016), have orthologues of TgDCX, proteins with DCX and P25 alpha together in the same molecule. Outside this group, this domain arrangement is only found in an early diverging metazoan (*Trichoplax adherens*, phylum Placozoa) (Orosz, 2009)

Proteins containing either the ~40 amino acid (aa) P25-alpha domain or the ~70 aa DCX domain are generally involved in interactions with microtubules. In vertebrates, proteins containing either of these domains are restricted to brain tissue (Gleeson et al., 1999; Liliom et al., 1999). In humans, mutations in the X-linked *doublecortin* gene, which codes for a protein containing tandem DCX domains, disturb neuronal migration to the cortex in the developing central nervous system, resulting in the double cortex syndrome in females and the more severe X-linked lissencephaly in males. Doublecortin, by binding to the groove between protofilaments in microtubules, is believed to play a role in preferentially stabilizing particular microtubule structures (Moores et al., 2004). P25-alpha domain proteins also modulate the structure of polymers formed from tubulin (Hlavanda et al., 2002). P25-alpha and DCX-domain proteins are thus candidates for specifying the unusual polymeric form of tubulin in the conoid fibers. In this report we show that loss of TgDCX radically disrupts the structure of the conoid. severely impairs host cell invasion, and slows growth.

## Results

### TgDCX gene structure and cloning

TgDCX was initially identified in the apical complex as one of ~170 proteins in a comparative proteomics analysis of conoid-enriched and conoid depleted fractions of *T. gondii* (Hu et al., 2006). The TgDCX gene was cloned from a *T. gondii* RH strain cDNA library. 5’ and 3’ cDNA ends were determined by rapid amplification of cDNA ends (RACE) experiments. Five (5’RACE) or four (3’RACE) clones were analyzed. Although there was clone to clone variation, the differences were small, <15 bp at each end. The 5’ end was mapped at (minus)532~(minus)513 bp from the putative translational start site of TgDCX and the 3’ end was mapped at 679~685 bp from the last base of the stop codon. The transcript from *T. gondii* is predicted to code for a 256 aa protein of 29.2 kDa. Comparison of gDNA and cDNA sequences indicated that there are 6 exons in this gene in *T gondii* (Figure 2), and gene models for most of the apicomplexans also predict 6 exons. The estimated molecular weights of TgDCX orthologues range from 19.1 kDa in *Trichoplax* to 33.3 kDa in *Cryptosporidium*. There is a well-conserved ~80aa DCX-domain (a brain-specific microtubule-interacting domain (Gleeson et al., 1999), part of the ubiquitin superfamily (Kim et al., 2003) near the C-terminus, which makes up slightly less than a third of the entire protein length in TgDCX. Another well-conserved region, rich in charged residues, is present in the middle, with similarity to a portion of TPPP/P25-alpha, a brain-specific tubulin polymerization promoting protein (Liliom et al., 1999; Hlavanda et al., 2002). The amino-terminal part is the least conserved.

**Figure 2.**
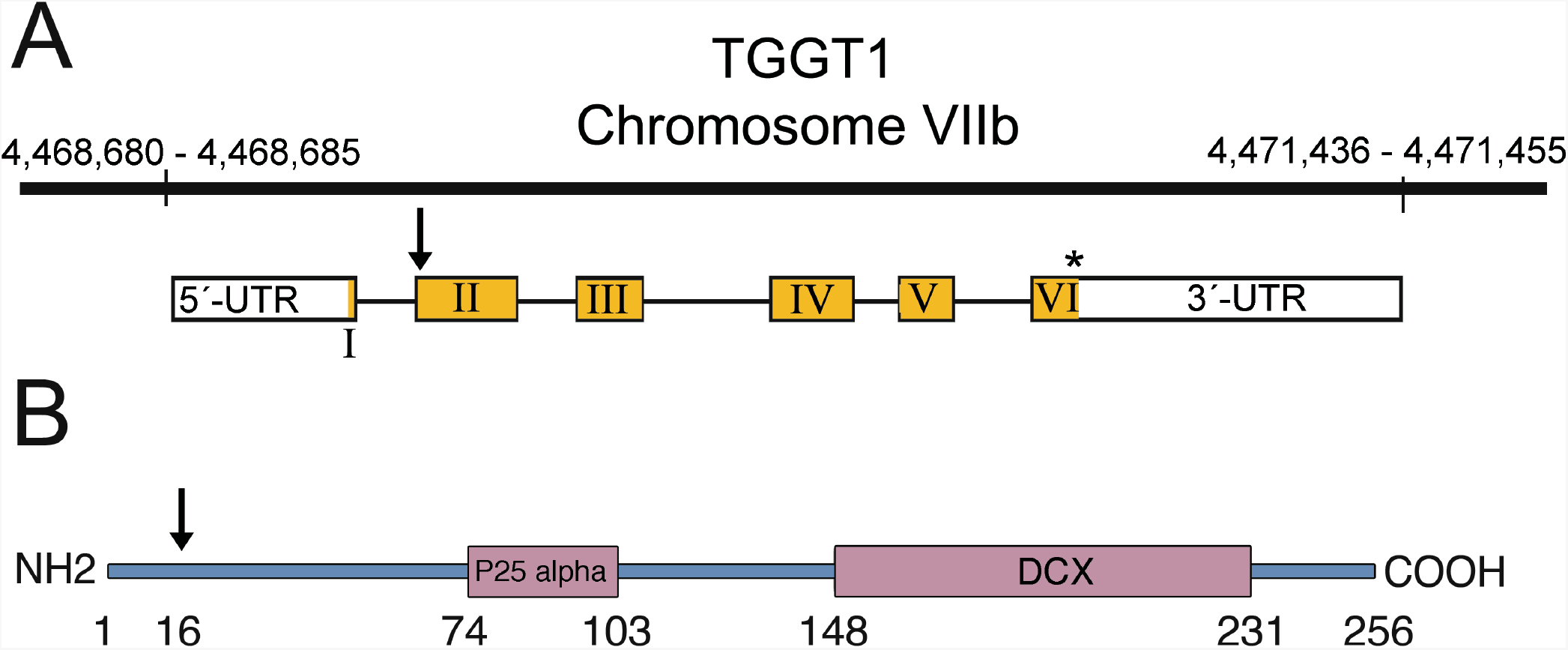
TgDCX gene (TGGT1-256030) A) Arrangement of TgDCX on TGGT1 Chromosome Vllb and structure of the mRNA. Non-coding regions are indicated by the white boxes. Exons are in yellow. The arrow indicates an alternative start codon. The stop codon is marked by an asterisk. B) Schematic of TgDCX protein. Regions with significant homology to known eukaryotic proteins are marked by the purple boxes. The arrow indicates an alternative translation initiation site 15 aa residues from the first methionine. Numbers indicate amino acid residues counting from the first methionine.

### Localization of FP tagged TgDCX in T. gondii by live-cell imaging

Stable transformants of *T. gondii* were created that expressed FP-tagged TgDCX either as randomly integrated extra copies, or by homologous recombination as a replacement for the endogenous gene under control of the native promoter (“DCX knock-in” parasites). Homologous recombinants expressing any of mCherry-TgDCX, TgDCX-mCherry, or TgDCX-mNeonGreen, all showed the same localization pattern, and their growth rate was indistinguishable from the parental line (RHΔHXΔku80). Fluorescence from FP-tagged TgDCX in adult parasites and developing daughters of the homologous recombinant lines is restricted to the conoid (Figure 3).

**Figure 3.**
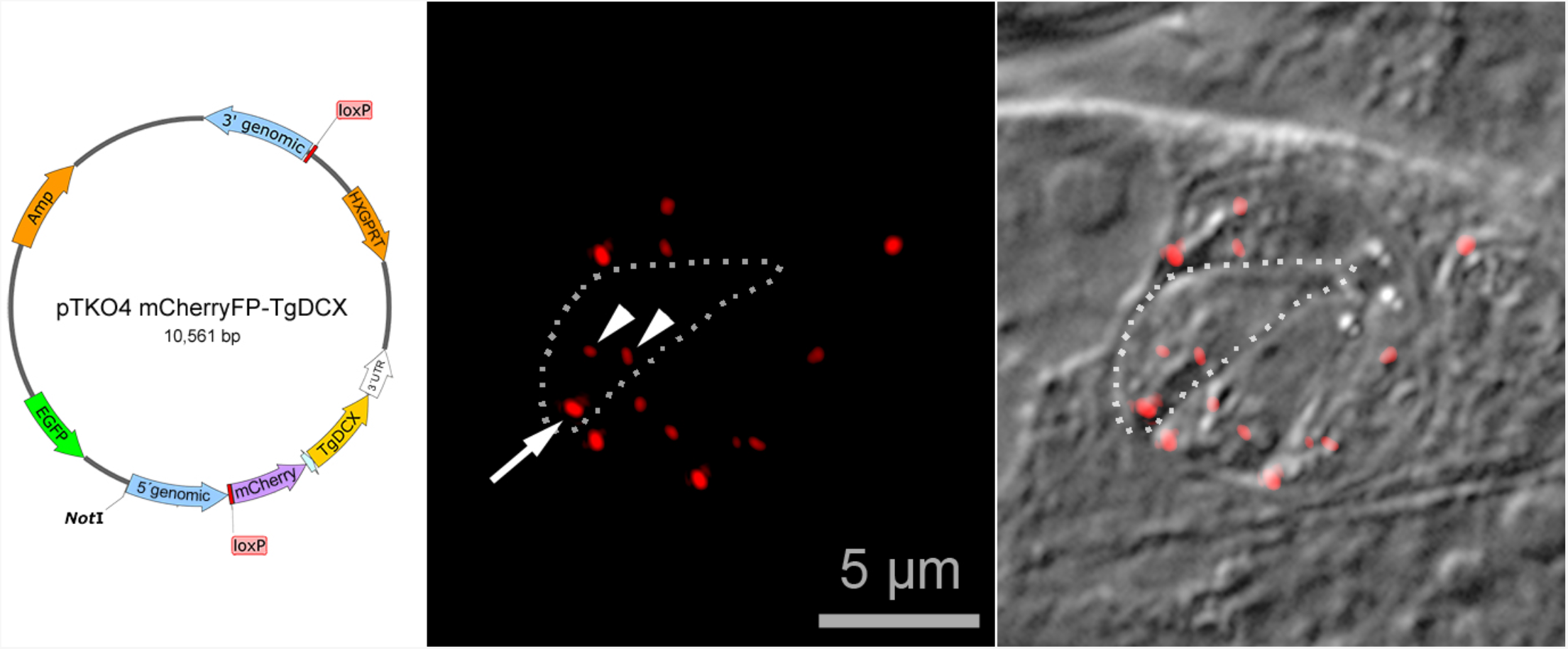
Left: Map of a TgDCX knock-in plasmid. Sandwiched between two loxP sites are a selection marker (HXGPRT expression cassette), the coding sequence for mCherryFP-tagged TgDCX, and the 3‘UTR from the T. gondii GRA2 gene. Flanking the loxP sites are 3’ and 5’ sequences, ~900 bp each, copied from T. gondii genomic DNA, that direct integration of the loxP sandwich into the genome by homologous integration at the TgDCX locus, replacing the entire TgDCX coding region, exons and introns. The plasmid is linearized by NotI cutting prior to electroporation into a ΔHXGPRT line of T. gondii. Integration of the knock-in plasmid at any site by a single crossover event preserves the EGFP expression cassette, allowing selection for true homologous recombinants. Integration at the DCX locus by homologous recombination removes the region of the plasmid not framed by the 3’ and 5’ homology regions. After homologous recombination, expressing Cre recombinase in knock-in parasites deletes all TgDCX and HXGPRT coding sequences from the genome. Right: Fluorescence and DIC images of mCherryFP-TgDCX knock-in parasites. The field of view includes one vacuole containing four parasites, and another vacuole with one parasite. The mCherryFP fluorescence is located in a single spot at the apical end of both adult (arrow) and developing daughter (arrowheads) parasites. One parasite of the four-parasite vacuole is enclosed by the dotted line.

### Estimating the number of molecules of TgDCX in the conoid

In the DCX-knock-in parasites, every molecule of TgDCX in the cell carries an identical fluorescent tag. Thus it should be possible to use the intensity of fluorescence in DCX-knock-in parasites to count the number of molecules of TgDCX in the conoid, a number whose value is required to formulate hypotheses about the mechanism of TgDCX stabilization of conoid fibers. In order to count by using fluorescence, some type of calibration is required, to provide an estimate of the amount of fluorescence detected from a single molecule of FP. Several different types of fluorescent calibration standard have been reported (for review, see (Verdaasdonk et al., 2014). We constructed a set of fluorescent standards by fusing various fluorescent proteins to the N-terminus of the E2 inner capsid glycoprotein of Sindbis virus, a small alphavirus, whose inner capsid has been shown to have T=4 icosahedral symmetry by 3D reconstruction from cryoEM images; that is, the inner capsid of each viral particle contains 240 copies of the FP-E2 fusion protein (Jose et al., 2015). Measurements of the fluorescence of conoids in the mCherry-TgDCX-knock-in parasites and mCherry-E2 Sindbis virions under the same illumination conditions showed that the conoids are 14–15 fold brighter than the virions, corresponding to ~3500 TgDCX molecules per conoid in the adult parasite.

For this estimate to be accurate, the photochemical properties of mCherryFP (i.e., extinction coefficient and quantum yield for fluorescence) when fused to TgDCX located in a conoid must be the same as when fused to glycoprotein E2 located in the nucleocapsid of the Sindbis virion. We cannot directly measure extinction coefficient and quantum yield for fluorescence for mCherryFP in the conoid, but we are able to measure the rate of photobleaching, which is determined by the extinction coefficient and quantum yield for photobleaching. Under the illumination conditions used for the brightness comparison, the rate constants and 95% confidence limits for photobleaching of mCherryFP in the Sindbis virion and in the *Toxoplasma* conoid are 0.0319 ± 0.0002 sec^−1^ and 0.0299 ± 0.0008 sec^−1^ respectively. This close agreement suggests that the fluorescent Sindbis virions are an appropriate fluorescent standard.

### TgDCX appears very early in daughter development

Earlier work with parasites expressing FP-tagged tubulin has shown that the initiation of the nascent daughter apical complex is marked by the appearance of a bright spot of tubulin distinct from the tubulin located in the spindle pole/spindle [(Hu, 2002; Hu, 2008; Nishi et al., 2008)]. With the advantage of structured illumination microscopy (SIM), that spot of tubulin can now be resolved into a 5-petaled flowerlike arrangement (Figure 4), as previously observed with a microtubule-associated protein (Liu et al., 2016). Co-expressing FP-tagged tubulin in the mCherry-TgDCX knock-in line reveals that in early daughter development, a spot of TgDCX appears in the nascent daughter apical cytoskeleton soon after tubulin, at the center of the 5-petaled flower (Figure 4).

**Figure 4.**
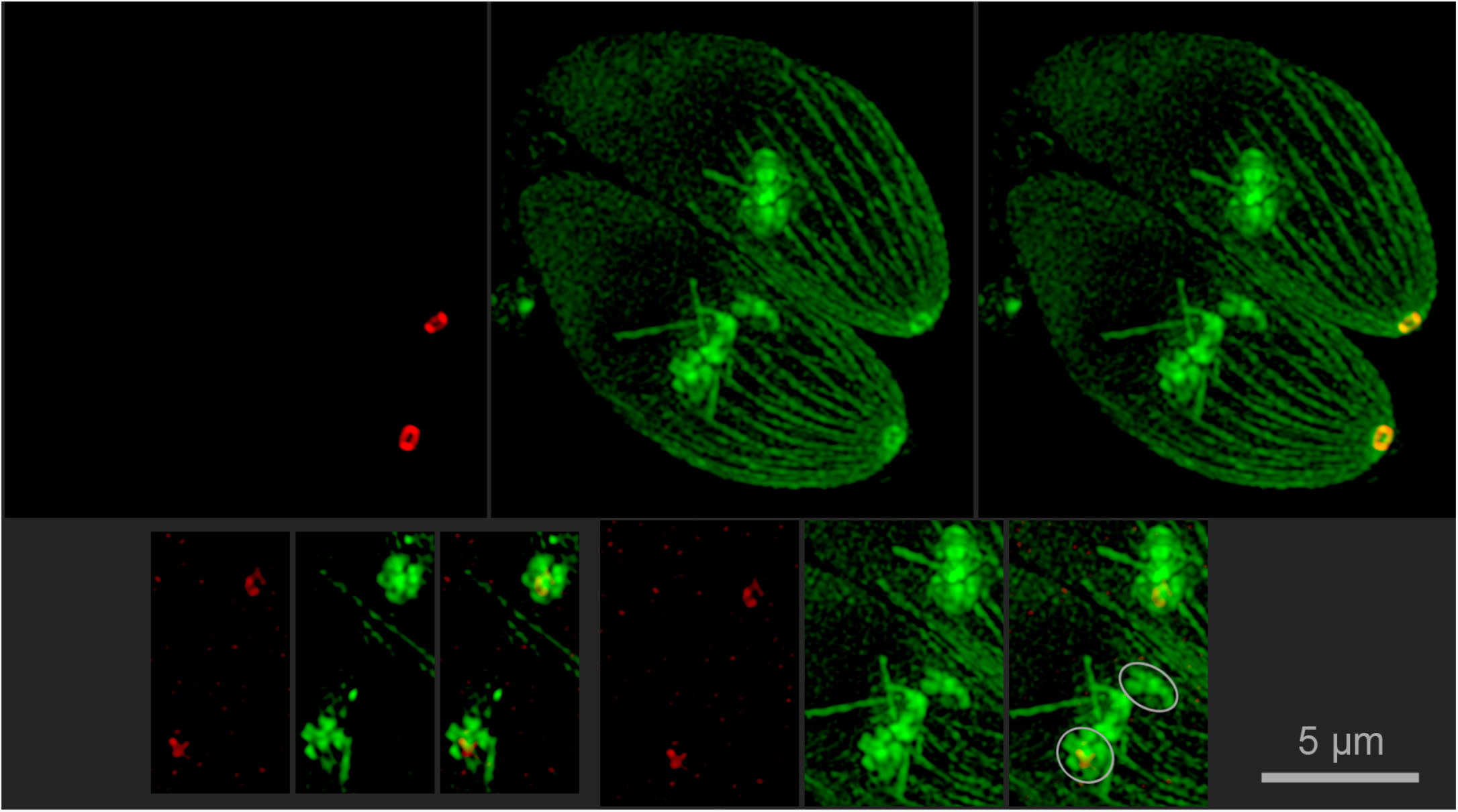
SIM images of mCherry-TgDCX (red) and NeonGreenFP-tubulin in TgDCX-knock-in parasites. Top row: a Z-projection of a 3D stack of images (25 slices) of a vacuole with two adult parasites at the initiation of daughter parasite formation. Left to right; mCherryFP-TgDCX, NeonGreenFP-tubulin, and overlay. Bottom row left: Single channel and overlay images from a single slice of the 3D stack showing the 5-petaled flower of tubulin that marks the beginning of daughter apical cytoskeleton formation. The slice happened to include one flower from each of the two adults. Bottom row right: Single channel and overlay images of the sum of 13 slices from the middle of the 3D stack showing the developing daughter apical cytoskeleton and the mitotic spindles in both adult cells. The 5-petaled flowers of the lower adult are indicated by the grey ellipses. One is viewed looking down the developing daughter apical-basal axis, the other is seen from the side. Note that the red channel of the images in the bottom row has been strongly contrast enhanced to make the small spot of mCherry-TgDCX visible. Its true intensity is ~1% of the adult mCherry-TgDCX spot; not visible in this display without contrast enhancement.

### Localization of TgDCX in T. gondii by antibody staining

An anti-TgDCX antibody was produced in rabbits by immunizing with recombinant TgDCX protein purified from bacteria. Two bands of unequal intensity were detected by Western blotting analysis of *T. gondii* whole cell lysate using this rabbit antibody (Figure 5). The estimated molecular masses of the two bands were 30.3 and 28.3 kDa, close to the theoretical molecular masses calculated based on the two different translation initiation sites (29.2 and 27.5 kDa). Immunofluorescent staining of *T. gondii* showed localization at the daughter and adult parasite apical complex (conoid) only (Figure 5). Immuno-EM labeling of extracellular parasites with rabbit anti-TgDCX and 1.4 nm gold particles with silver enhancement shows TgDCX only in the tubulin-containing conoid fibers (Figure 6). No other structures are labeled.

**Figure 5.**
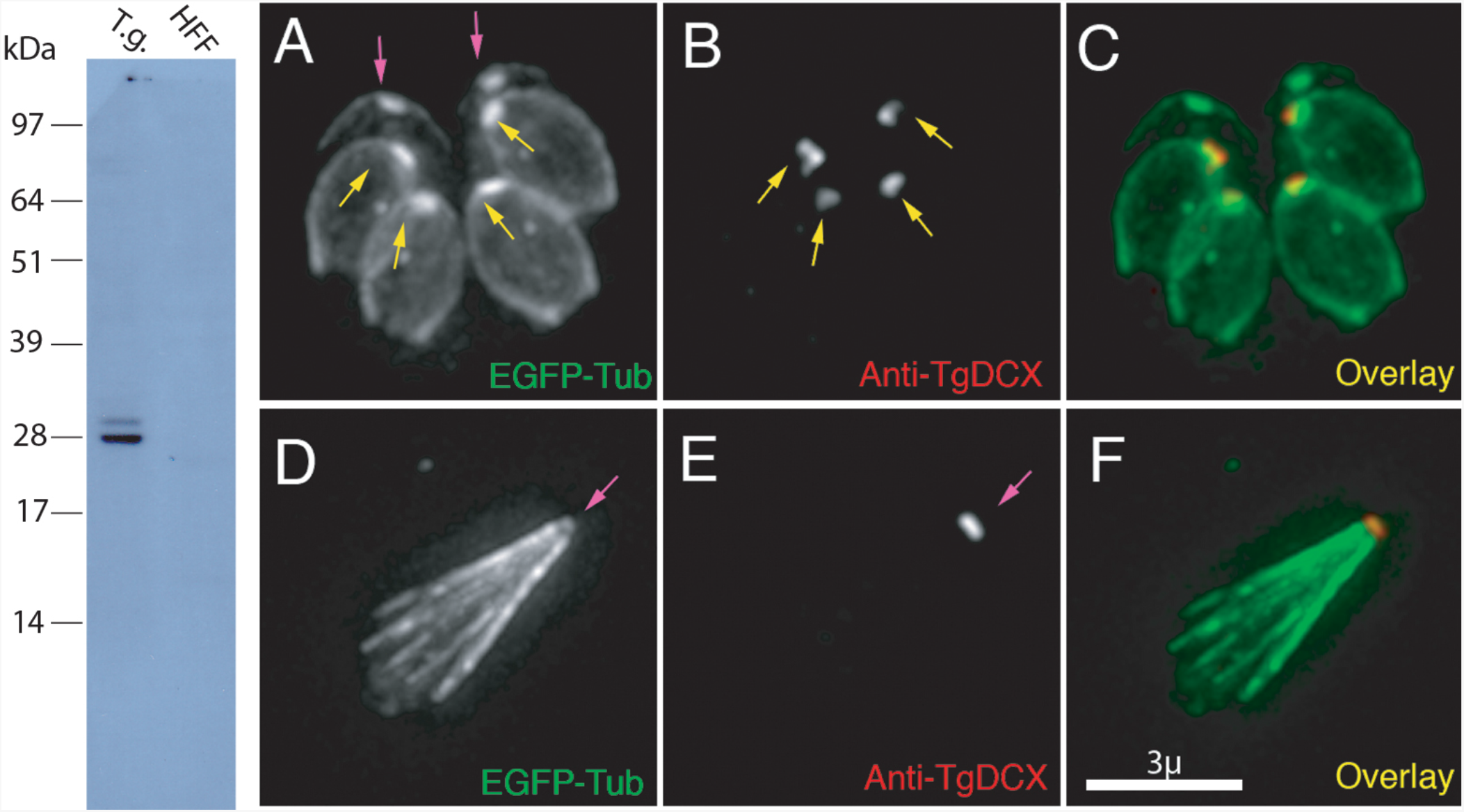
Left: Western blot with rabbit antibody raised against bacterially expressed recombinant TgDCX. Lane 1 (T.g.) Whole-cell lysate of RH parasites, 10 µg protein, 7 × 10^6^ cells. Lane 2 (HFF) 10 µg protein from whole cell lysate of uninfected host cells (human foreskin fibroblasts) Right: Images of a transgenic RH line expressing EGFP-β3-tubulin (green) and stained with rabbit anti-TgDCX antibody (red). Top row: Intracellular parasites fixed and permeabilized with methanol and briefly treated with 10 mM deoxycholate. With this protocol, or using non-ionic detergent permeabilization, the developing daughter conoids are labeled, but antibody does not penetrate the adult conoid. Anti-GFP antibody behaves similarly - in parasites expressing TgDCX-EGFP, daughter conoids can be stained with a GFP-antibody, but adult conoids are not labeled, even though the adult conoids are brightly fluorescent from the DCX-EGFP they contain (data not shown). Bottom row: using longer deoxycholate treatment to permeabilize/extract extracellular parasites allows the antibody to penetrate the adult conoid, but the architecture of the parasite is mostly disrupted (cf. Figure 6).

**Figure 6.**
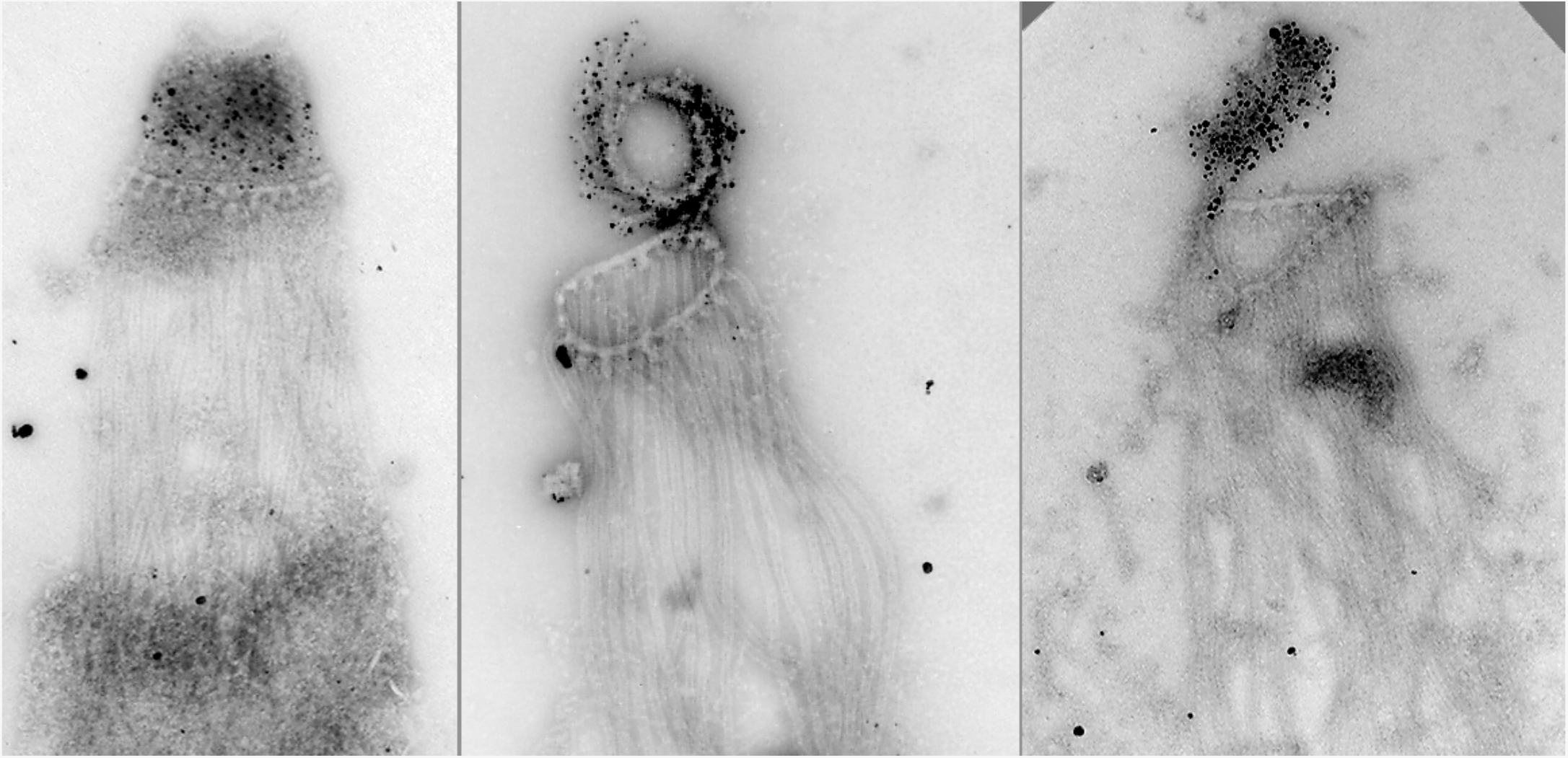
EM images of deoxycholate extracted RH parasites labeled with antiTgDCX antibody, 1.4 nm gold-secondary antibody, and silver enhanced. The silver/gold deposits are confined to the conoid. Cortical microtubules are not labeled. In the middle image, the conoid is partially uncoiled, revealing individual conoid fibers with attached gold/silver deposits.

### Creation and phenotype of TgDCX null mutant lines of Toxoplasma gondii

The Cre-recombinase method described earlier [(Heaslip et al., 2010; Liu et al., 2016)] was used in conjunction with the three homologous recombinant lines described above to delete *TgDCX* from the genome. A population of slow-growing but viable parasites was obtained (“DCX-knockout” parasites), from which ten clonal lines were established, two clones derived from the mCherryFP-TgDCX knockin line, four from the TgDCX-mCherryFP knockin, and four from the TgDCX-mNeonGreenFP knockin. The loss of TgDCX protein was confirmed by the disappearance of the fluorescence due to FP-tagged TgDCX. The deletion of the TgDCX coding sequence from the genome was verified by Southern blot (Figure 7).

**Figure 7.**
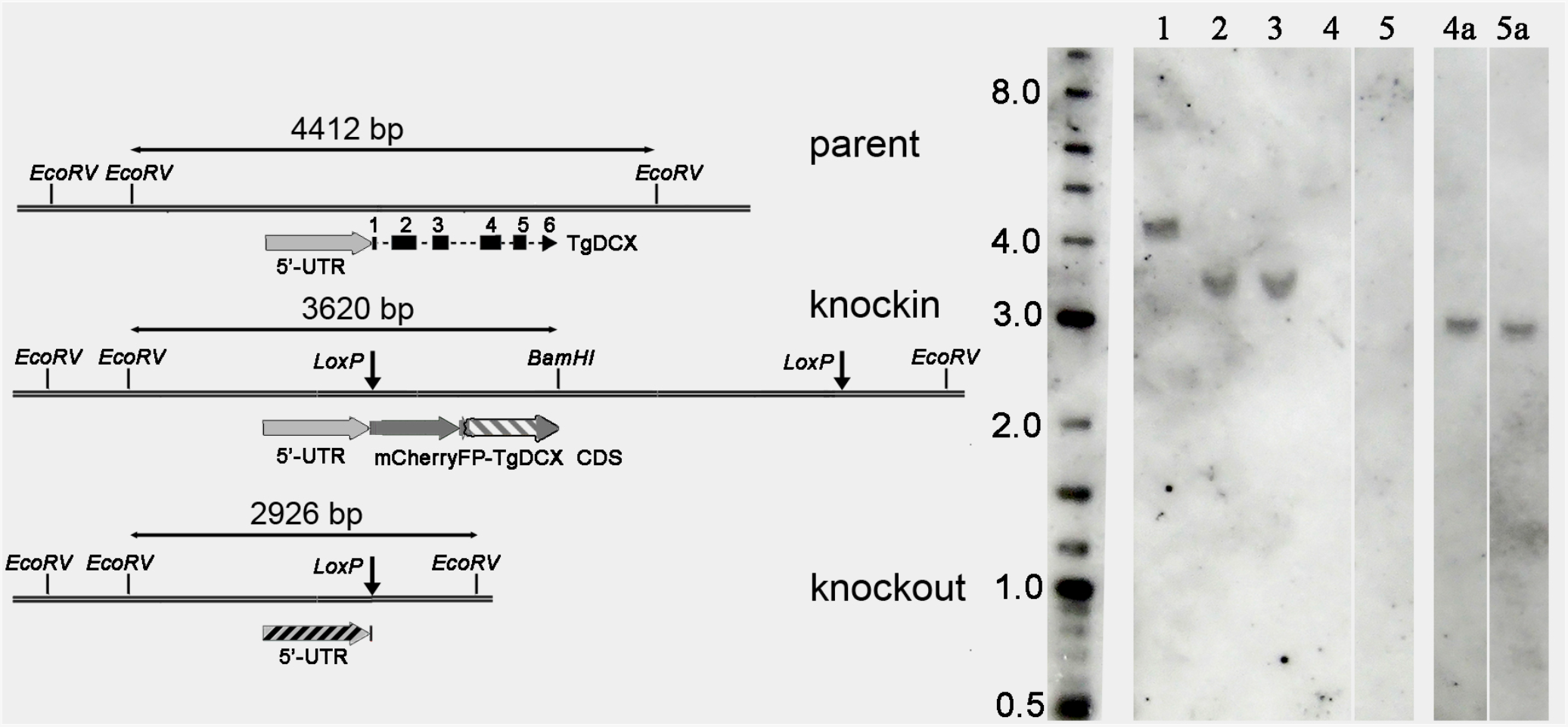
Southern blot with T. gondii genomic DNA. Left side: Map of the TgDCX genomic region in parental (RHΔHXΔKu80), knock-in (FP-TgDCX or TgDCX-FP), and knockout (ΔTgDCX) parasites. The sizes of the BamHI-EcoRV or EcoRV-EcoRV fragments containing the hybridization targets are indicated. Sequence regions included in the probes are indicated by the striped regions; exon 2–5 probe, grey-white stripes in knock-in map; 5’-UTR probe, grey-black stripes in the knockout map. The 5’-UTR region is identical in the three parasite genomes. Right side: size marker (length in kbp); Lanes 1–5: hybridized with a probe against exons 2–5 of TgDCX CDS. lane 1, RHΔHXΔKu80 parasites, expected size 4412 bp; lane 2 and 3, mCherryFP-TgDCX and TgDCX-NeonGreenFP knock-in parasites, expected size 3620 bp; lane 4 and 5, clones of knockout parasites derived from mCherryFP-TgDCX and TgDCX-NeonGreenFP knock-in parasites respectively, no band expected. Lanes 4a and 5a are lanes 4 and 5 respectively of the original blot after it was stripped and re-hybridized with a probe against the 5’-UTR region of the TgDCX gene. expected size 2926 bp.

The growth of these TgDCX-null lines was much slower than the parental line. Under conditions where the parental line showed a doubling time (see *Methods*) of ~7.5 hours, the DCX-knockout line doubled in ~17 hours. However, with continued culture some adaptation occurred, with the growth rate stabilizing after ~ 4 weeks of serial passage in host cells. In two independent clones of the DCX-knockout parasite line maintained in continuous culture for several months, the doubling time had decreased to ~13 hours. In a plaque assay performed after ~10 weeks of continuous culture, the DCX-knockout parasite line formed approximately one-fourth as many plaques as the parental line, and the plaques were about one-third the size (Figure 8). Complementation of the null mutant with an exogenous copy of the TgDCX CDS, tagged with EGFP and expressed under control of the *Toxoplasma* α-tubulin promoter, restored the number of plaques to approximately the same as the mCherryFP-TgDCX knock-in line (Table I).

**Figure 8.**
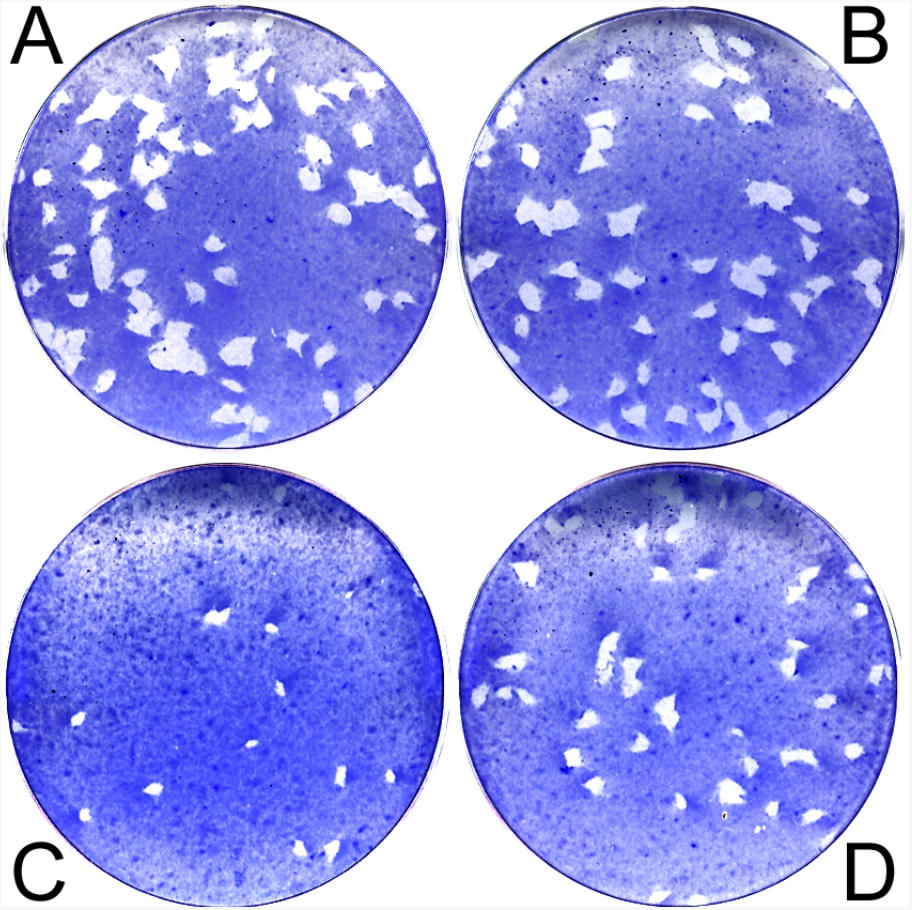
Plaque assay. A, parental (RHΔHXΔKu80); B, knock-in (mCherryFP-TgDCX); C, knockout (ΔTgDCX); D, knockout parasites complemented with TgDCX-EGFP expressed under control of the T. gondii α-tubulin promoter.

**Table I.** Host cell lysis and invasion by T. gondii lines. The average number and size of plaques (± std. error) formed in three replicate experiments are tabulated. Cytolytic efficiency (“cyt. eff.”) is defined as the ratio of the total area lysed by a line to the total area lysed by the parental line, expressed as a percentage.

|  | plaque assay |  |  | invasion assay |  |
| --- | --- | --- | --- | --- | --- |
|  | # of plaques | plaque size (mm<sup>2</sup>) | cyt. eff. (%) | # invaded | % of parental |
| RHΔHXΔku80 | 61 ± 7 | 21 ± 1 | 100 | 2435 ± 84 | 100 |
| mCherryFP-DCX | 58 ± 3 | 20 ± 0.3 | 92 | 1199 ± 85 | 49 |
| ΔDCX | 18 ± 1 | 8 ± 1 | 12 | 628 ± 43 | 26 |
| ΔDCX+EGFP-DCX | 52 ± 6 | 14 ± 0.4 | 55 | 1244 ± 79 | 51 |

To further characterize the growth defect seen in TgDCX-knockout parasites, we assayed host-cell invasion (Carey et al., 2004) by the parental, knock-in, knock-out, and “complemented” knockout parasite lines (Table I). In this assay, monolayer cultures of host cells are exposed to a suspension of parasites for a short time, then rinsed with fresh medium, fixed and labeled with an antibody against a *Toxoplasma* surface antigen and secondary antibody coupled to a red fluorophore. Parasites that had adhered to host cells but had not yet invaded are accessible to the antibody and therefore labeled with red fluorescence. Parasites that had invaded into host cells are not accessible to the antibody and so are not stained. The host cells are then permeabilized with detergent, and the antibody labeling is repeated, this time using a secondary antibody tagged with a green fluorophore. Thus when the assay is complete, intracellular parasites are labeled green, but those parasites on the outside of the host cell are labeled red, enabling easy discrimination between completed invasion events and merely adhering to the host cell surface.

Replacing the endogenous TgDCX with mCherryFP-TgDCX (DCX knock-in parasites) reduced host cell invasion in our assay by approximately two-fold. Complete loss of TgDCX caused a further two-fold reduction. Adding back a copy of the TgDCX CDS tagged with EGFP, randomly inserted in the genome, restored host cell invasion to the same level as the mCherryFP-TgDCX-knock-in parasites. Note that whereas the native DCX promoter drives expression in the mCherryFP-TgDCX-knock-in parasites, in the TgDCX-EGFP complemented knockout parasite, expression is driven by a mis-matched promoter (*T. gondii* α-tubulin gene).

### Loss of TgDCX is associated with a loss of tubulin in the apical complex

Extension of the conoid/apical complex induced by calcium-ionophore treatment in the DCX-knockout parasites, as judged by phase-contrast light microscopy, appeared indistinguishable from that observed in the DCX-knock-in lines or in parental wild-type parasites (RH or RHΔHXΔKu80) (data not shown). However, examination of DCX-knockout parasites in which all tubulin containing structures were made visible by transforming with a plasmid driving expression of NeonGreenFP α-tubulin revealed that the apical complex region of the ΔTgDCX parasites was abnormal, with a much reduced or sometimes complete absence of the normally observed bright apical spot of fluorescence due to tubulin in the conoid. Figure 9 shows SIM images of NeonGreenFP α-tubulin fluorescence in wild-type and knockout parasites. Fluorescence arising from the cortical microtubules in the adult and daughter knockout parasites appeared normal, as did the intra-conoid microtubules, mitotic spindle, spindle pole, and centrioles. The only noticeable difference is in the conoid, which is clearly resolved in slices through the apical region of parasites whose long axis happened to be oriented perpendicular to the optical axis. In such slices (enlargements on the right side of Figure 9), the conoid is observed as a bright apical ring or short cylinder in normal parasites, but is greatly diminished or absent in DCX-knockout parasites. Interestingly, at least two other proteins normally found in the apical complex, AKMT (Heaslip et al., 2011) and TgCAP2 (Leveque et al., 2016) are not obviously affected by the loss of TgDCX, as judged by immunofluorescent localization in knockout-parasites by wide-field epifluorescence (data not shown).

**Figure 9.**
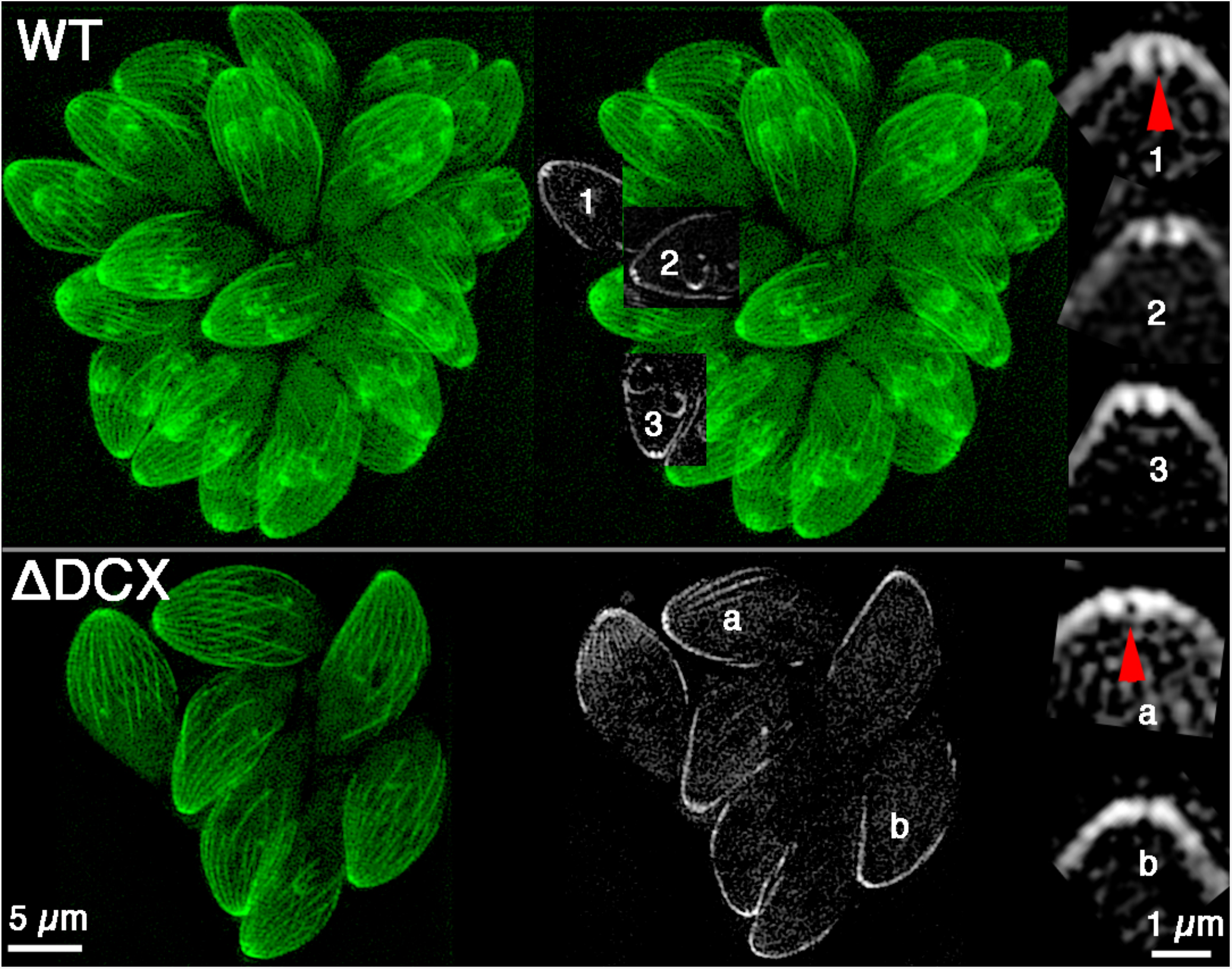
SIM images of parental and ΔTgDCX knockout parasites expressing NeonGreen-β-tubulin. Left column: Z-projections of the entire 3D stacks of images. Middle column: a single slice (lower row) or pieces of 3 different slices superimposed on the Z-projection (upper row). Right column: 4× magnified portions of single slices through the conoid regions of 3 (upper) or 2 (lower) parasites. Arrowheadss indicate the conoid region of the corresponding insets.

In interpreting sets of SIM images such as Figure 9 that contain data from different specimens, it is necessary to keep in mind the relationship between the brightness in the displayed images and the fluorescence intensity in the actual specimens. The brightness of the enlarged images on the right of Figure 9 were scaled individually to utilize the full 8-bit range of the display, because unfortunately there seems currently to be no satisfactory way of objectively normalizing the relative intensities in SIM images of different specimens. In the parental parasites, the conoid spot is much brighter than anything else in the selected subregion. Consequently, the arch of the cortical microtubules is dimmer than the conoid in the displayed images of those parasites. In contrast, in the knockout parasites, the arch of the cortical microtubules is the brightest feature, because the spot of tubulin in the conoid region is much reduced. For that reason, the arch of cortical microtubules appears much brighter in the knockout parasites than in the parental parasites in the display, whereas in reality they probably have approximately the same fluorescence intensity.

### Deletion of TgDCX causes severe structural defects in the conoid

Electron microscopy of negatively stained TgDCX-knockout parasites revealed gross structural abnormalities in the conoid. Figure 10 shows a comparison of conoids from normal wild-type parasites with conoids from both the DCX-knock-in and the DCX-knockout lines. The parasites were treated with calcium ionophore to induce conoid extension, and briefly with mild detergent to permit visualization of the conoid by negative staining. The conoid of the mCherryFP-TgDCX-knock-in parasites, from which the knockout parasites were derived, is indistinguishable from wild-type at this resolution, but the TgDCX-knockout is abnormal. The wild-type is characterized by a prominent basket-weave stripe pattern, arising from superposition of the front and back sides of the spiral of 14 conoid fibers [(Hu et al., 2002)]. This stripe pattern is weaker or sometimes absent, less regular, and less extensive in conoids from the knockout lines. Overall, the knockout conoid is narrower, shorter, and less rectangular in profile than the wild-type. Whereas the wild-type conoid looks much the same in every cell, the knock-out conoid is quite variable in appearance. Figure 11 shows a montage of conoid images from randomly selected cells of TgDCX-knockout parasites to illustrate the range of variation. Some are less disrupted than others, but in an extended survey of several hundred TgDCX-knockout parasites, we have not observed one that appears normal. In some cases, the overall shape of the conoid in the knockout parasites resembles a partially extended conoid in normal parasites, but in detail they are not alike. Figure 12 shows that in normal parasites, even with a retracted conoid, the conoid fibers are still prominent and appear well-ordered.

**Figure 10.**
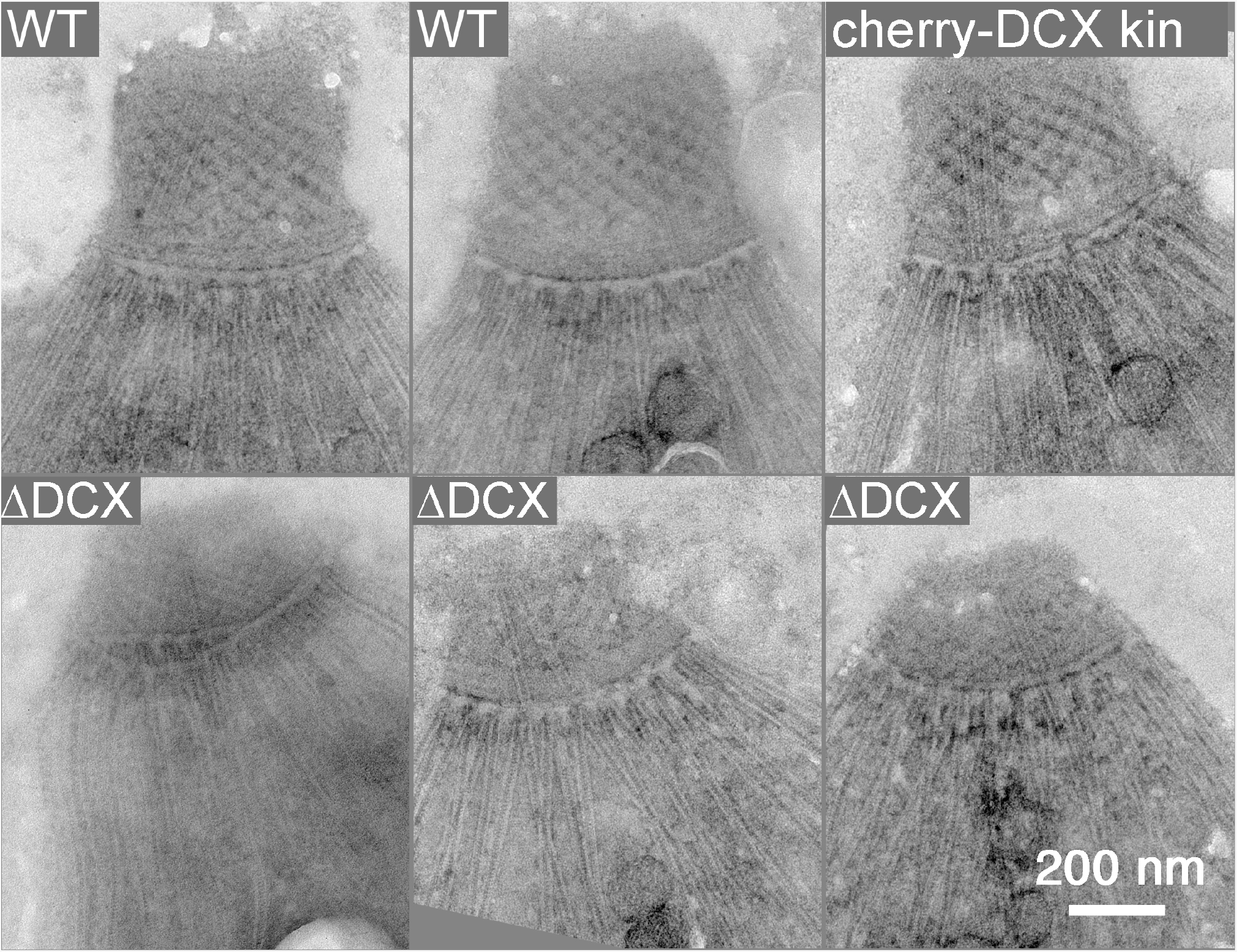
EM images of the conoid region of negatively stained T. gondii. WT, parental line RHΔHXΔku80, two examples; cherry-DCX kin, mCherryFP-TgDCX knock-in parasites; ΔDCX, TgDCX knockout parasites, 3 examples.

**Figure 11.**
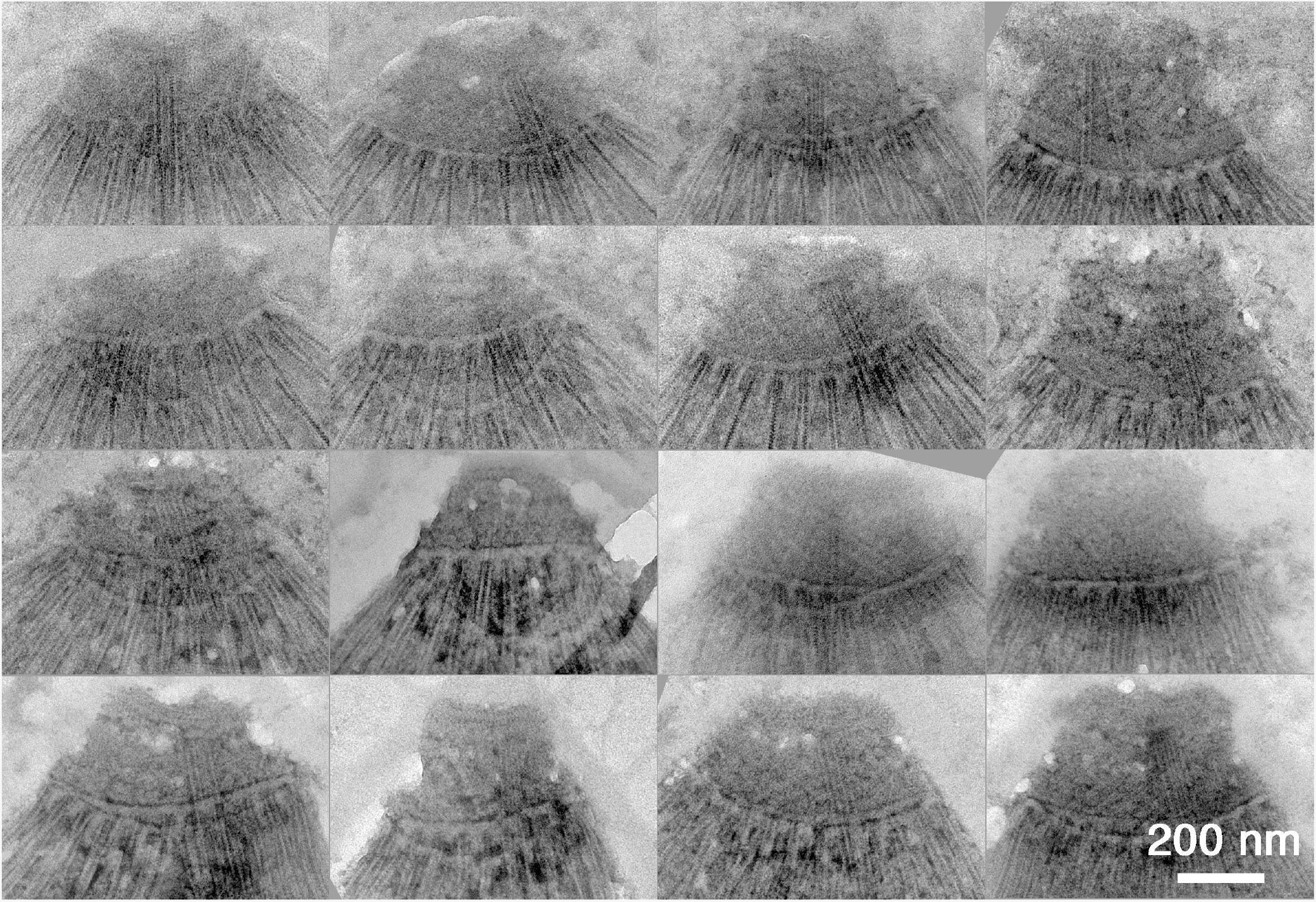
Montage of EM images of the conoid region of 16 randomly chosen negatively stained TgDCX knockout parasites.

**Figure 12.**
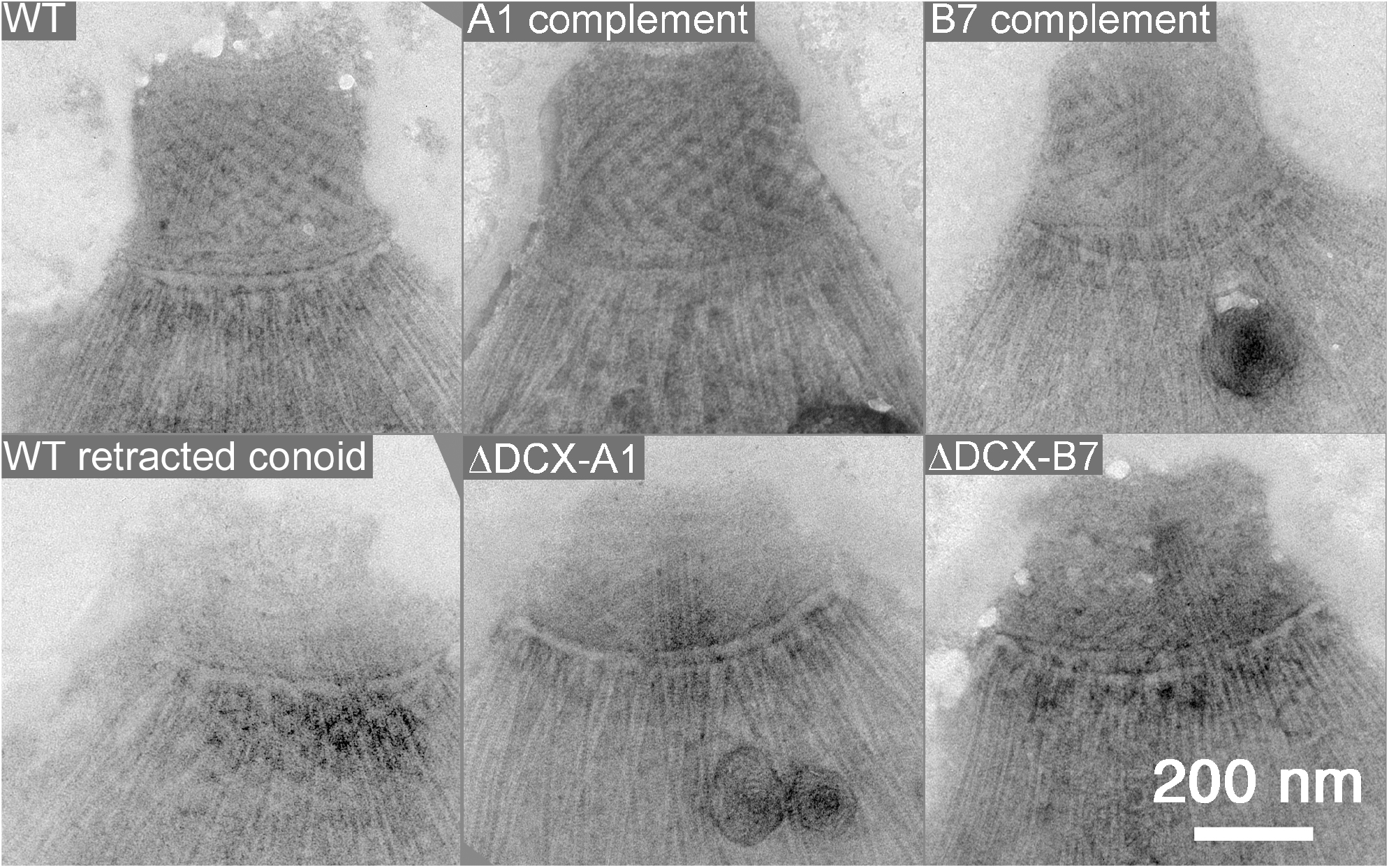
EM images of the conoid region from wild-type T. gondii, extended and retracted, two clones of TgDCX knockout parasites, A1 and B7, and the same two clones after transforming with a plasmid driving expression of TgDCX-EGFP (A1 complement, B7 complement).

To confirm that the observed structural defect is indeed due to the loss of DCX protein, we transformed TgDCX-knockout parasites with a plasmid containing the TgDCX CDS fused to EGFP, with 5’-UTR (promoter) from the *T. gondii* α-tubulin gene, and 3’-UTR from the *T. gondii* dihydrofolate reductase gene. By fluorescence microscopy, the TgDCX-EGFP is localized to a bright spot at the apical end of these “complemented knockout” parasites (data not shown). Figure 12 shows that the structural defect in the conoid of the TgDCX-knockout parasites is completely reversed by expression of the TgDCX CDS.

The conoid fibers contain tubulin dimers that are essentially identical in primary amino acid sequence to the tubulin dimers in the cortical microtubules, intraconoid microtubules, and other tubulin containing structures in the cell. Tubulin expressed from extra copies of the α-tubulin gene or any of the three β-tubulin isoforms (Hu et al., 2003), even human tubulin expressed in *T. gondii* (Nagayasu et al., 2006) is incorporated promiscuously into all tubulin containing structures, and yet TgDCX is normally found only on the conoid fibers, and removal of TgDCX affects only the conoid fibers, not any other tubulin containing structure. This surprising independence extends in both directions. Figure 13 shows images from a recently described (Liu et al., 2016) parasite line lacking three microtubule associated proteins (TLAP2, SPM1, TLAP3). In this triple knockout line, the cortical and intraconoid microtubules are absent in the TX-100 extracted parasites, but the conoid fibers are completely normal. The exquisite precision revealed by the differential localization and independent effects of these tubulin binding proteins is remarkable.

**Figure 13.**
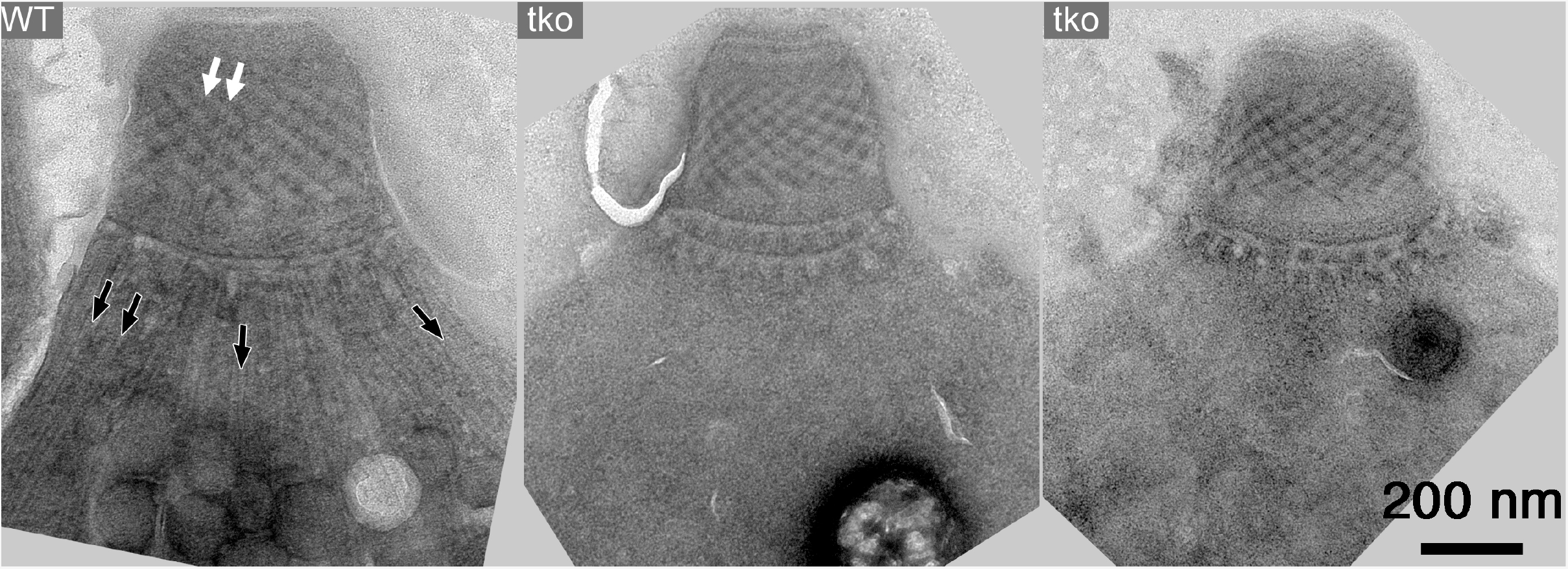
EM images of the conoid region from wild-type T. gondii (“WT”) and Δtlap2Δspm1Δtlap3 parasites, which lack three microtubule associated proteins (Liu et al., 2016). The 22 cortical (black arrows) and 2 intraconoid (white arrows) microtubules are missing in the triple knockout parasites, but the conoid fibers are intact.

## Discussion

The common feature of all members of the phylum Apicomplexa is the presence of an apical complex, an assemblage of cytoskeletal elements and secretory organelles at one end of the cell (Figure 1). Although the exact structures of the apical complex vary among members, it is believed to serve important functions in both invasion of host cells and in replication of the parasites. Many members of the phylum have in their apical complex a distinctive organelle, the “conoid”, a ~380nm diameter motile organelle, consisting (in *T. gondii*) of 14 spirally-wound conoid fibers, which are non-microtubule polymers built from tubulin subunits. It has been hypothesized that the apical complex is structurally related to the flagellum, based on the fact that a number of flagellar components such as dynein light chain and SAS6-like protein are found in the apical complex in *T. gondii* (Hu et al., 2006; de Leon et al., 2013). Further, organisms in the sister clades that possess a pseudoconoid invariably have flagella rooted close to the apical complex (Portman and Slapeta, 2014). *T. gondii* also forms a flagellum during its sexual stage when the parasite differentiates into microgametes in the epithelial cells of the small intestine of a cat. However, the structural relationship between the apical complex and the flagellum remains unknown in *T. gondii* because it is difficult to obtain microgametes. It is also not known whether the apical complex and the flagella in chromerids and perkinsids indeed share common structural origins or they are simply unrelated structures that happen to be located in the vicinity of each other.

The conoids of *T. gondii* and its close relatives in the Apicomplexa differ from those of more distant relatives not only in being closed cones rather than half-closed cones or flat ribbons, but also in the size of the organelle and consequent curvature of the fibers composing it. In *T. gondii*, the conoid fibers are bent into an arc with a radius of curvature of ~ 200nm. The rigidity of microtubules in cells is increased to some extent by binding of associated proteins (Kurachi et al., 1995), but even when completely bare, ordinary microtubules break when bent into arcs with radii less than ~500 nm (Amos and Amos, 1991). Thus a transition from the large conoids of other Alveolates to the compact conoid of Apicomplexans may require special modifications of the conoid fibers to decrease their flexural rigidity. Among those modifications is presumably the change from a tubular (ordinary microtubules) to a convex ribbon (*T. gondii* conoid fibers) polymer structure. However, such a structural modification is not possible using tubulin alone, because the tubulin dimers in a convex ribbon cannot all be in equivalent local environments. Whereas it is possible to make a cylindrical (helical) polymer of protofilaments of tubulin dimers in which every dimer is equivalent, that is not possible with a convex ribbon. The energy to support this sort of non-equivalence must be supplied in some form, most commonly in the form of the binding energy of non-tubulin associated proteins. If that source of extra energy were to be removed, for instance by removal of the associated protein, the polymer with non-equivalent subunits would become less stable. This seems a reasonable explanation for the destabilization of the conoid fibers and loss of tubulin from the conoid in the TgDCX knockout parasites. This explanation predicts that tubulin polymerized in the presence of TgDCX would assemble into some structure other than a canonical microtubule, perhaps a curved convex ribbon. Unfortunately it has not yet been possible to test this prediction because bacterially expressed TgDCX remains in solution only under denaturing conditions, but planned future experiments using a baculovirus expression system may provide the opportunity.

By comparing the fluorescent intensity of conoids in an mCherryFP-TgDCX-knock-in line of *T. gondii* with that of an alphavirus particle containing 240 copies of mCherryFP, we estimated that a conoid contains approximately 3500 TgDCX molecules. The knock-in parasite invades host cells slightly less well than the wild-type parasite, raising the possibility that the FP tag hinders DCX incorporation into the conoid, so 3500 may be a lower bound for the true number of DCX molecules in the wild-type conoid. The 14 conoid fibers are approximately 400 nm long and contain 9–10 protofilaments, with tubulin dimer spacing of 8 nm along the protofilament (the same spacing as in a canonical microtubule). Therefore there are 6300 - 7000 tubulin dimers per conoid, or no less than one DCX molecule per two tubulin dimers. If DCX is located uniformly along all conoid fibers (consistent with, though certainly not proved by, Figure 6), they would thus be spaced at 16 nm intervals. In terms of protein mass, and therefore of scattering power in cryoEM, the contribution from DCX (~ 30 kDa) must be ~15% of that from tubulin (dimer ~ 100kDa), sufficient to account for the observed 16 nm periodicity seen in Fourier transforms of cryoEM images of conoid fibers (Hu et al., 2002).

Orthologues of TgDCX are present in all sequenced apicomplexan genomes, even in those lacking a recognizable conoid. Its ubiquitous presence suggests that the acquisition of the DCX gene occurred early in apicomplexan evolution. The continued existence of the TgDCX orthologues in apicomplexans that lack distinctive conoid structures, such as *Cryptosporidium*, poses interesting questions. Part of the answer may lie in differences between life-cycle stages. For instance, although it seems that haematozoea species (haemosporidians and piroplasms) have lost the conoid from their apical complexes in meroozite and sporozoite stages, the ookinete stages in some species still retain conoids (Desser, 1970; Patra and Vinetz, 2012). This may indicate that the loss of conoid in haematozoea is a relatively recent event, which suggests that it would be worth looking carefully in those species for some remnant or radically modified form of conoid. Localization studies of DCX in those seemingly conoid-less species, particularly in their ookinete stages, could provide valuable insight into this interesting question.

Deletion of TgDCX significantly decreases the rate of host cell invasion. In our assay, in which parasites are presented with host cell targets for a limited time (one hour), wild-type parasites invade approximately four times faster than TgDCX-knockout parasites. In the less exacting setting of a plaque assay or in continuous culture, where the time allowed for invasion is essentially unlimited, the difference between wild-type and knockout parasites is reduced, resulting in an overall difference in apparent growth rate of only 2-fold. However, both of these situations are artificial and not representative of the situation faced by the parasite attempting to survive and spread in an intact animal with a functioning immune system. In that situation, the parasite is protected when enclosed in its intracellular vacuole, but highly vulnerable when extracellular. Thus there must be enormous selection pressure acting to make re-invasion after egress as fast as possible.

Our experiments do not provide any insight into exactly how TgDCX facilitates host cell invasion. It might be that its stabilization of the conoid fibers provides some beneficial mechanical advantage for host cell penetration, or alternatively, render the intra-conoid passageway a better conduit for secretion from the parasite of the effector molecules known to be important during invasion. Further work will be required to distinguish between these possibilities, but the availability of the knock-in, knockout, and complemented knockout parasite lines we have established provide the tools needed to make the distinction experimentally.

## Methods

### Culture, harvest, and transformation of Toxoplasma gondii

*T. gondii* tachyzoites were used in all experiments, and grown in monolayers of human foreskin fibroblast (HFF) cells (Roos et al., 1994). Parasites were harvested from culture supernatant by centrifugation at 3600 × g for 1 min when the monolayer was ~80% lysed. For purification, the parasite suspension was passed through a 3 µm Nucleopore filter (Whatman 110612), centrifuged, and resuspended in DPBS at ~4 × 10^8^/mL. Transformation of *T. gondii* tachyzoites was carried out as previously described (Heaslip et al., 2009) using 30–40 µg of plasmid DNA in “cytomix” buffer (120 mM KCl; 0.15 mM CaCl_2_; 10 mM KH_2_PO_4_ / K_2_HPO_4_; 25 mM HEPES; 2 mM EGTA; 5 mM MgCl_2_, 2 mM K_2_ATP, 5 mM glutathione, pH 7.6).

### *Cloning of TgDCX* (All PCR primers are listed in Table 2)

#### 5’RACE

At the time TgDCX was cloned (Nagayasu et al., 2006), the various *T. gondii* genome annotations contained conflicting gene models for a hypothetical protein, varying in length from 131 to 1109 amino acids. A template cDNA was synthesized from *T. gondii* RH strain total RNA by SuperScriptII RT (Invitrogen, Carlsbad, CA) using reverse primer AS1, complementary to nucleotides 705~726 counted from the then putative translational start of the hypothetical protein, 750–771 from the start of the ultimately cloned TgDCX (all subsequent numbers refer to the current TgDCX gene model), followed by dA-tailing using dATP and terminal transferase. The product was subjected to 3 rounds of hemi-nested PCR. For the first round, oligo(dT)-anchor primers (SAS1, SAS2, and SAS3), an anchor primer without oligo(dT) (SAS4) and a gene specific primer AS2 (position 382–402) were used. For the second and third rounds, the same anchor primer (SAS4) was used in combination with either AS3 (for second round, position 167–186) or AS4 (for third round, position 82–101). The final PCR products were purified on a 1.0% agarose gel, then cloned into the vector pCR2.1 TOPO (Invitrogen). Five clones were sequenced and compared with the genomic DNA sequence obtained from the public database (www.toxodb.org).

**Table II:**
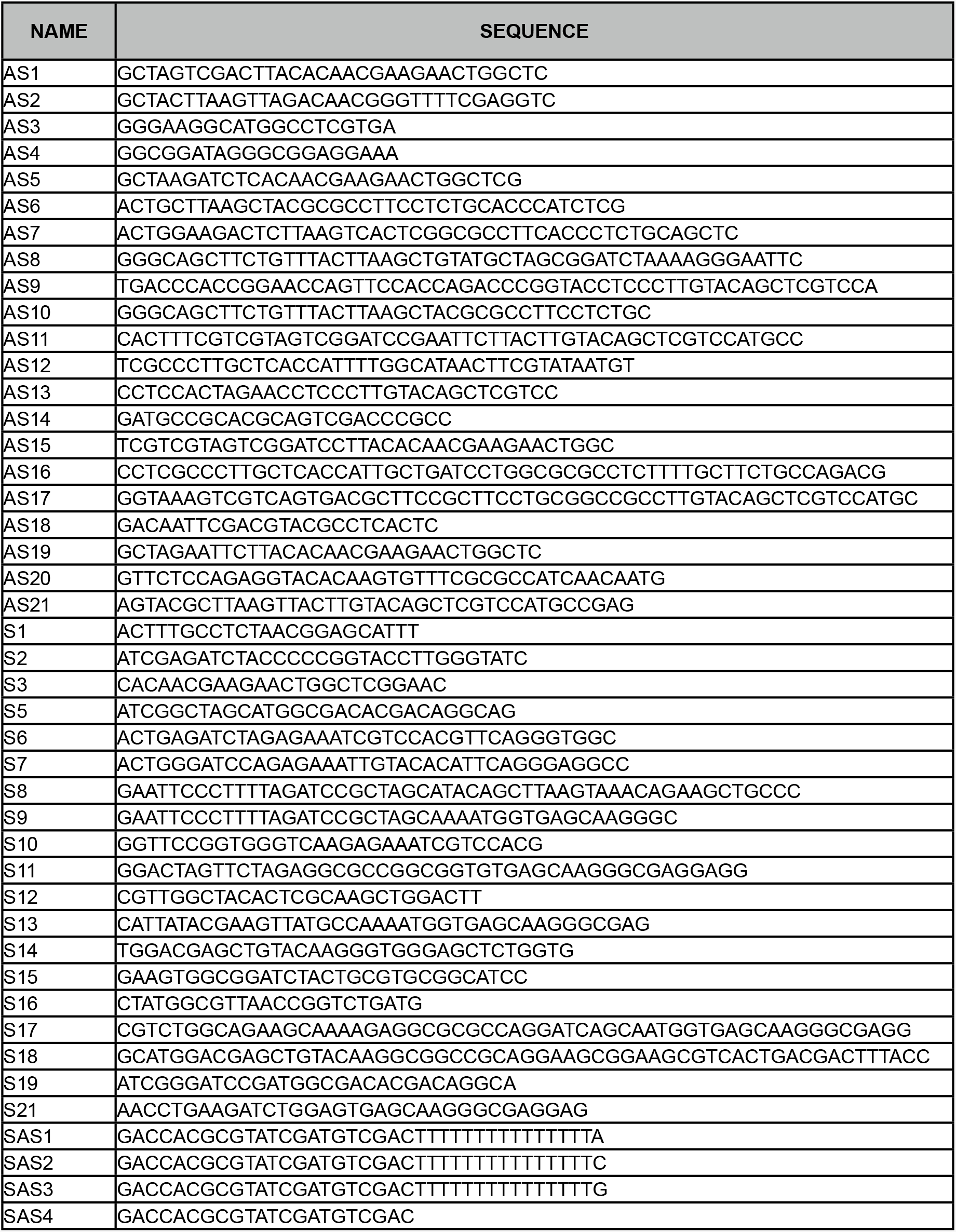
Oligonucleotides used in this study

#### 3’RACE

3’ RACE ready cDNA was synthesized using *T. gondii* total RNA and the oligo(dT) anchor primers (SAS1/ SAS2/SAS3) by SuperScriptII RT. Three rounds of hemi-nested PCR were performed using the synthesized cDNA, forward primers S1 (first round, position (minus)194 to (minus)172), S2 (second round, position 403–423) then S3 (third round, position 745~768) in combination with the anchor primer (SAS4). The final PCR products were gel-purified, cloned then sequenced as for the 5’RACE products.

### *Plasmid construction* (All PCR primers are listed in Table 2)

After construction, plasmids were used to transform chemically competent TOP10 cells by heat shock, or electrocompetent DH5α cells (NEB C2989) by electroporation. Plasmid DNA was isolated by standard procedures and the constructions were verified by DNA sequencing.

ptub-TgDCX-EGFP is a derivative of the plasmid ptub-H2b-YFP (Hu et al., 2004). Converting from ptub-H2b-YFP to ptub-TgDCX-EGFP entailed removing the H2b sequence between the NheI and BglII sites and replacing it with the coding region of TgDCX extending from the second methionine to the stop codon, obtained by RT-PCR using tachyzoite total RNA and primer pair S5/AS5. YFP was swapped with EGFP via the BglII and AflII restriction sites and a coding sequence of EGFP PCR amplified using primers S21 and AS21. In the final expressed product, TgDCX is coupled to the N-terminus of EGFP (lacking its initial methionine) via the 3-residue amino acid sequence RSG. Another version of ptub-TgDCX-EGFP was also constructed, with amino acid substitution Y219H in the TgDCX CDS (numbered according to position in the full length protein), corresponding to a doublecortin mutation giving rise to lissencephaly in humans (human Y125H). By light microscopy, the localization of these two forms of TgDCX-EGFP did not differ significantly from each other or from wild-type. Both were effective in complementing the structural defects of the conoid in TgDCX knockout parasites by EM analysis and in complementing its growth defect by plaque assay.

ptub-EGFP-TgtubB1 and ptub-EGFP-TgtubB3 are both based on the ptub vector backbone. In both cases, the EGFP coding sequence is inserted between the NheI and BglII sites, the tubulin coding sequence lacking the initial methionine is inserted between the BglII and AflII sites. The β1-tubulin coding sequence was obtained by RT-PCR using tachyzoite total RNA and primer pair S6/AS6, cut with BglII and AflII and ligated into the vector backbone. The β3-tubulin coding sequence was amplified by RT-PCR from tachyzoite total RNA with primer pair S7/AS7, cloned into vector PCR4-TOPO (Invitrogen), cut with BamHI and BbsI to give BglII-AflII compatible ends, and ligated into the BglII-AflII cut vector backbone. In the expressed protein product the EGFP is in both cases linked to the tubulin by the 5 amino acid sequence SGLRS.

ptub-mNeonGreenFP-TgtubB1 was derived from the plasmid ptubg. Plasmid ptubg was created from ptub-EGFP-TgtubB1 by a 2 component assembly using the NEBuilder^®^ HiFi DNA Assembly reagents (NEB E2621S), following the manufacturer’s recommended protocol. Component 1 was prepared from ptub-EGFP-TgtubB1 by cutting out the 2088 bp NheI-AflII fragment. Component 2 was prepared by mixing equal volumes of oligonucleotides S8 and AS8, 10 µM each, heating to 97°C for 2 min, and cooling slowly to room temperature. Plasmid ptubg-mNeonGreenFP-TgtubB1 was constructed by a 3 component NEBuilder^®^ HiFi assembly. Component 1 was prepared by cutting out the 12 bp stuffer from ptubg using NheI and AflII. Component 2 was prepared by PCR amplifying the mNeonGreenFP coding sequence from pmNeonGreen-N1 (Shaner et al., 2013) using primers S9 and AS9. Component 3 was prepared by PCR amplification of the *T. gondii* β1-tubulin coding sequence from plasmid ptub-EGFP-TgtubB1 with primers S10 and AS10. In the expressed protein product, mNeonGreenFP is coupled to the N-terminus of β1-tubulin (lacking its initial methionine residue) by the 13 aa flexible linker G-(GTGSG)2-GS.

The plasmid pTKO4-TgDCX-mCherryFP was constructed in a backbone (“pTKO4”) designed for replacement of genes in *T. gondii* with mCherryFP-tagged versions of the gene by homologous recombination. This backbone was derived from pTKO2_II (Heaslip et al., 2010) by several modifications. (1) The EGFP coding sequence was replaced by a synthesized version of EGFP in which restriction sites for AflII, BpmI, MfeI, NcoI, and NsiI were removed by silent mutagenesis. (2) The three MCS sequences were replaced (see below). (3) mCherryFP was added, positioned so that it can be fused to the C-terminus of the target CDS.

MCS1 in pTKO2_II, which is used for insertion of a piece of the genomic DNA upstream of the target gene just in front of a LoxP site, encoded sites for NotI, PstI, KpnI, XhoI, and EcoRV. This was replaced with a synthesized sequence encoding sites for SgrDI, BsiWI, NotI, AflII, HinDIII, EcoRI, and BglII.

MCS2 in pTKO2_II, which is used for insertion of the coding region of the target gene on the other side of the LoxP site, encoded sites for EcoRI, BglII, PmeI, AsiSI, RsrII, and StuI. This was replaced with a synthesized sequence encoding sites for PacI, NsiI, SphI, AsiSI, NheI, MfeI, PmeI, SpeI, XbaI, and MreI. Also inserted downstream of this new MCS2 was a synthesized sequence for mCherryFP minus the initial methionine codon. The last “G” of the MreI site of MCS2 is also the first base of the valine codon (GTG) in the N-terminal amino acid sequence (MVSKGEE) of mCherryFP. The sequence TAAGGATCC was also added on the 3’-end of the mCherryFP coding sequence (stop codon plus BamHI site).

MCS3 in pTKO2_II, which is used for insertion of a piece of the genomic DNA downstream of the target gene, is separated from MCS2 by the mCherryFP coding sequence, a portion of the 3’-UTR of TgGRA2 (serves as 3’-UTR for the target gene after homologous recombination), the expression cassette for TgHXGPRT (for selection of transfected cells with 80 µM mycophenolic acid plus 330 µM xanthine), and a second LoxP site to allow knockout of the target gene by Cre recombinase (Heaslip et al., 2010). MCS3 encoded sites for HinDIII, NheI, HpaI, ApaI, and PspOMI. This was replaced with a synthesized sequence encoding sites for BclI, SacI, AscI, XhoI, AbSI, KpnI, SmaI, XmaI, FseI, SfiI, ApaI, and PspOMI.

To construct pTKO4-TgDCX-mCherryFP, the 887 bp sequence of *T. gondii* RH genomic DNA immediately preceding the methionine initiation codon of TgDCX was inserted into MCS1 between the AflII and EcoRI sites. The sequence between the last base of the EcoRI site of MCSI and the last base of the PmeI site of MCS2 was replaced with a synthesized piece that included a LoxP site (34 bp), a *T. gondii* consensus Kozak sequence (6 bp), and the complete TgDCX CDS (768 bp). With this insertion, the TgDCX CDS is then fused to the N-terminus of mCherryFP (lacking initial ATG) by a 9aa linker PGTSSRGAG. Finally, a synthesized sequence extending from the last 8 bp of the TgDCX Exon VI in *T. gondii* RH genomic DNA, through the stop codon and the first 871 bp of the downstream genomic region, flanked by 5’-AscI and 3’-AbsI recognition sites was inserted using the AscI and AbsI sites of MCS3.

To construct pTKO4-TgDCX-mNeonGreenFP, mCherryFP was removed from pTKO4-TgDCX-mCherryFP by cutting with BamHI and MreI, and replaced a by PCR amplified copy of the NeonGreenFP (Shaner et al., 2013) coding sequence (primers S11 and AS11) using NEBuilder^®^ HiFi DNA Assembly. In the final construct, the TgDCX CDS is fused to the N-terminus of mNeonGreenFP (lacking initial ATG) by a 10aa linker PGTSSRGAGG.

pTKO4-mCherryFP-TgDCX was constructed from pTKO4-TgDCX-mCherryFP by a 5-component assembly using the NEBuilder^®^ HiFi assembly reagents. Component 1was obtained from pTKO4-TgDCX-mCherryFP by removing the 1757 bp BamHI-NheI fragment. Component 2 was obtained by PCR copying the piece of TgDCX 5’-UTR and adjacent LoxP site from pTKO4-TgDCX-mCherryFP using primers AS12 and S12. Component 3 was obtained by PCR amplifying an mCherryFP coding sequence (from which XcmI, HinCII, PvuII, BbSI, BtgI, PstI, and MscI sites had previously been removed by silent mutagenesis) using primers S13 and AS13. Component 4 was obtained by PCR amplifying a 69 bp synthetic linker sequence using primers S14 and AS14. Component 5 was obtained by PCR copying the complete TgDCX CDS plus stop codon, minus the initial ATG codon, from pTKO4-TgDCX-mCherryFP using primers AS15 and S15. In the final construct, mCherryFP is coupled to the N-terminus of TgDCX (minus the initial methionine) via a 23 bp flexible linker, GGSS-(GGS)6-T.

To construct TE12-mCherryFP, a plasmid (kindly provided by Dr. Suchetana Mukhopadhyay, Indiana University, Bloomington IN) encoding the complete genome of recombinant Sindbis virus TE12 (Lustig et al., 1988) was cut with BsiWI and HpaI, removing the 3462 bp region coding for proteins C, E3, E2, 6K, and part of E1. This segment was replaced with a 4206 bp piece, flanked by the same restriction enzyme sites, containing the same coding sequences plus mCherryFP fused in frame between E3 and E2. The 4206 bp piece was produced by overlap PCR from three fragments. Fragment 1 (1758 bp) was amplified from the TE12 plasmid, position 6910-8634, with primers S16 and AS16. Fragment 2 (785 bp) was amplified from a synthesized mCherryFP coding sequence (from which XcmI, HinCII, PvuII, BbSI, BtgI, PstI, and MscI sites had been removed by silent mutagenesis) using primers S17 and AS17. Fragment 3 (1804 bp) was amplified from TE12 position 8629-10396 with primers S18 and AS18. These three fragments were purified, mixed, and used as template for PCR with primers S16 and AS18. The 4206 bp product was purified, cut with BsiWI and HpaI, and ligated into the TE12 vector. In the expressed protein product (before cleavage by furin), E3 is coupled to the N-terminus of mCherryFP with the 5 aa linker GAPGSA, and mCherryFP is coupled to the N-terminus of E2 with the 6 aa linker AAAGSG. TE12-mNeonGreenFP was prepared in the same way, except fragment 2 for the overlap PCR was obtained by amplifying the mNeonGreen coding sequence with primers S17 and AS17.

### Fluorescent Sindbis virus production

Viral RNA (+)-strand was prepared from the TE12-FP plasmids by *in vitro* transcription with SP6 RNA polymerase (Rice et al., 1987). Plasmid DNA (100–200 ng) was linearized by cutting with PvuI, precipitated with ethanol, and re-dissolved in 20 µL of 40mM TrisCl (pH 7.9 at room temperature), 10mM MgCl_2_, 2mM spermidine, 10 mM DTT, 2mM each of rATP, rCTP, rGTP, and rUTP, 0.5 mM 3’-O-Me-m7G(5’)ppp(5’)G RNA cap structure analog (New England Biolabs S1411), 1X RNAsecure™ reagent (Life Technologies AM7005), and incubated at 60 °C for 10 min. After cooling to room temperature, 20 units of SUPERase• In™ RNase inhibitor and 20 units of SP6 RNA polymerase (New England Biolabs M0207) were added, the mixture was incubated at 39 °C for 2 hours, and the entire reaction mix was immediately used for transfection.

Each *in vitro* transcribed RNA was transfected into Vero or BHK-21 cells in a 24-well plate using Effectene (Qiagen 301425) following the manufacturer’s protocol, with a total volume of 180 µL of serum free media per well. After overnight incubation at 37 °C in a CO2 incubator, 0.8 mL of culture medium was added and incubation continued for an additional 24 hours. Plaques of virus infected cells were easily visible by fluorescence microscopy using a low power (4× or 10×) objective. 48–72 hours after transfection, the culture supernatant was collected, centrifuged at 16,000×g for 5 min to remove cell debris, then stored in 0.1 mL aliquots frozen at −80 °C. Prior to use for large-scale virus production, an aliquot was thawed and the concentration of infective virus particles determined by plaque assay (Hernandez et al., 2010).

### Creating mCherryFP-TgDCX, TgDCX-mCherryFP, and TgDCX mNeonGreenFP knock-in parasites

Plasmids pTKO4-TgDCX-mCherryFP, pTKO4-TgDCX-mNeonGreenFP, and pTKO4-mCherryFP-TgDCX (40 µg each) were linearized by cutting with NotI, and separately introduced into extracellular RHAHXΔku80 parasites by electroporation. After 24 hours, selection for parasites containing a functional HXGPRT derived from the introduced plasmid using 80 µM mycophenolic acid and 330 µM xanthine in the culture medium. After 10 days of drug selection a stable population was obtained, from which 27 individual non-GFP expressing (i.e., potentially homologous recombinants) clones were isolated by limiting dilution. Two of the candidate knock-in clones were eliminated by PCR screening for the intron-containing genomic version of TgDCX. Southern blotting was used to confirm that the remaining 25 were true homologous replacements.

### Creating a TgDCX knockout parasite

The pmin-Cre-eGFP_Gra-mCherry plasmid (Liu et al., 2016) was introduced into extracellular mCherryFP-TgDCX, TgDCX-mCherryFP, and TgDCX-mNeonGreenFP knockin parasites by electroporation. This plasmid drives expression of both Cre recombinase and mCherryFP. The mCherryFP is diffusely cytoplasmic and much brighter than the small dot of mCherryFP-TgDCX at the conoid, making it easy to recognize transfected parasites. Three cytoplasmic mCherry-positive parasites were sorted by FACS into each well of 96 well plates containing culture medium plus 0.5 mM 6-thioxanthine to select against parasites containing a functional HXGPRT. After plaques were visible, the wells containing single plagues were consolidated into a new 96-well plate and grown with no drug selection. Clones were screened for the absence of fluorescence. 10 “dark” clones were expanded and used to isolate genomic DNA. The absence of the TgDCX coding region was confirmed by Southern blotting.

### ptub-TgDCX-EGFP complemented parasites

To generate the complemented lines, ptub-TgDCX-EGFP plasmids were electroporated into TgDCX knockout parasites. The populations were then subjected to chloramphenicol selection for ~ one month and taken off drug selection for ~ 2 weeks before the growth of the populations was compared with the knockout, knock-in, and RHΔHXΔku80 parasites using plaque and invasion assays as shown in Figure 8 and Table 1.

### Recombinant TgDCX protein

The coding region of TgDCX extending from the second methionine to the stop codon, corresponding in size to the major band seen in Western blots of *T. gondii* whole cell extract (Figure 5), was obtained by RT-PCR with RH tachyzoite total RNA and primer pairs S19 and AS19, then cloned into the BamHI-EcoRI site of pRSET-B vector (Invitrogen), resulting in a fusion of 33 amino acids (hexahistidine-tag, T7 gene10 leader sequence, and Xpress epitope tag) to the N-terminus of the protein. The recombinant protein was produced in BL21 codon plus (DE3) RIL *Escherichia coli* (Stratagene, La Jolla, CA), and purified using Talon metal affinity resin (Clontech) under denaturing condition. Under non-denaturing conditions, the protein was found only in insoluble inclusion bodies. Extensive trials were undertaken in an attempt to prepare re-folded protein that is soluble in physiological buffers, without success.

### Antibody production

Polyclonal antiserum against the recombinant fusion protein was produced in rabbits by Cocalico Biologicals (Reamstown, PA). Although in some respects the results obtained with this antiserum were quite variable, in the one indispensable characteristic it proved reliable: in immunofluorescence, the only structure that it consistently labeled was the conoid, the same pattern seen with FP-tagged TgDCX, and in Western blots it always stained the recombinant or endogenous TgDCX bands. Under most conditions, in Western blots the antiserum also reacted with some higher molecular weight bands, even in extracts of host cells that were not infected with *T. gondii*. The reactivity against those extra bands could be attenuated or removed by several methods, including affinity purification, pre-absorption against fixed (Toxo-free) host cell monolayers, and pre-absorption against BSA or other unrelated proteins bound to nitrocellulose membrane. The recipes given below are those which were most successful in removing the non-specific reactivity.

Anti-TgDCX was affinity purified from the antiserum using the recombinant protein bound to nitrocellulose as follows. 160µg of recombinant TgDCX was run on a polyacrylamide gel and transferred onto a nitrocellulose membrane. The area of membrane which had the recombinant protein band was visualized by reversibly staining with Ponceau S, cut into small pieces, and incubated with 1mL of the crude antiserum against TgDCX for 2 hours at room temperature. The membrane pieces were washed twice with 0.1% Tween 20 / PBS (3 × 5 min). Bound antibodies were eluted in 900µL of 100mM glycine (pH2.5). The solution was neutralized with 100 µL of 1M Tris (pH8.0).

The affinity purified antibody solution was then subjected to the following pre-adsorption procedure to remove residual non-specific reactivity. HFF cells grown in a 75cm^2^ flask were rinsed with PBS then fixed with cold methanol for 5 min at −20°C. The cells were re-hydrated with PBS 3 × 5 min, blocked with 3% BSA, 0.01% Tween 20 / PBS for 1 hour at room temperature then washed once with PBS. 30uL of the affinity purified antibody solution was diluted 1:100 in PBS then incubated with the cells for 2 hours at room temperature. The pre-adsorbed antibody solution was collected from the flask and sodium azide was added to a final concentration of 0.02% for preservation at 4°C. This residual non-specific reactivity is removed equally well by pre-absorption with BSA or other proteins bound to nitrocellulose membrane.

### Western blotting analysis

Whole cell extract from 7 × 10^6^ parasites or 10 µg protein from uninfected HFF cells were mixed with NuPage sample buffer (Invitrogen) containing a final concentration of 10 mM DTT and run on 4%-12% NuPage Bis-Tris gel (Invitrogen). The separated proteins were transferred to a nitrocellulose or PVDF membrane. The membrane was blocked in 5% nonfat dry milk in PBS, incubated with the rabbit anti-TgDCX antibody diluted 1:5000, or pre-absorbed antibody diluted to 1:100, in blocking buffer + 0.1% Tween 20 for 1 hour at room temperature and washed in PBS-T (PBS + 0.1% Tween 20). The membrane was then incubated with horseradish-peroxidase-conjugated anti-rabbit IgG (Amersham) diluted 1:10000 in blocking buffer + 0.1% Tween 20 for 1 hour at room temperature, washed with PBS-T, then PBS. Chemiluminescent detection was performed using ECL-plus (Amersham).

### Immunofluorescence assays

HFF cells grown in 35mm glass bottom dishes (P35G-1.5-14-C or P35G-1.5-20-C, MatTek corporation) were infected with wild-type or transgenic tachyzoites expressing EGFP-Tgβ3-tubulin or TgDCX-EGFP. About 16 hours post-inoculation, cells were fixed with 0.5% formaldehyde in PBS for 5 min at room temperature, rinsed with PBS twice, re-fixed / permeabilized with cold methanol for 5 minutes at −20°C. After rinsing twice with PBS, cells were further extracted with 10mM deoxycholate in water for 5 minutes at room temperature with gentle rocking. Samples were rinsed with PBS twice then blocked with 3% BSA, 0.01% Tween 20 / PBS, followed by incubation with the rabbit anti-TgDCX for 1 hour at room temperature. Cells were washed with 0.01% Tween 20 / PBS (3× 5min) then incubated with goat anti-rabbit IgG conjugated with Alexa Fluor 555 (Molecular Probes, diluted 1:500 in blocking buffer), for 1 hour at room temperature. Cells were washed with 0.01% Tween 20 / PBS, 3 × 5min then mounted using ProLong anti-fade reagent (P7481, Molecular Probes).

Antibodies penetrate into adult conoids very poorly, making detection of antigens in adult conoids unreliable with typical immunostaining procedures, including the protocol above, which incorporates brief treatment with 10 mM deoxycholate. To circumvent this problem, a different procedure was developed using extracellular tachyzoites and more vigorous deoxycholate treatment. Freshly lysed-out parasites were harvested and passed through a 3 µm pore filter (Whatman, 111112). Cells were washed once with calcium saline (20mM HEPES, 2.7mM KCl, 138mM NaCl and 5mM CaCl2), pelleted again, resuspended in 200µL of warm conoid extension buffer (2µM A23187 in calcium saline) and incubated for 10 minutes at room temperature. The cell suspension was transferred to a glass bottom dish and the cells were allowed to settle. The supernatant was carefully removed and replaced with 500µL of extraction buffer (10mM deoxycholate in water supplemented with DNaseI (Invitrogen, 18068-015) then incubated with 15 minutes at room temperature. Cells were further extracted with 2mL of the extraction buffer without DNaseI for an additional hour with gentle rocking. Blocking, primary and secondary antibody reactions were performed similarly to the first immunostaining procedure described above.

### Immunoelectron microscopy

Immunogold labeling used essentially the second immunostaining procedure described above. Extracellular tachyzoites of wild-type RH strain were passed through a 3µm pore filter, washed once with calcium saline, pelleted and resuspended in 30µL of the warm conoid extension buffer, then incubated for 10 minutes at room temperature. Parasites were adsorbed onto a carbon-film-coated nickel grid for 10 minutes, incubated with 30µL of extraction buffer (10mM deoxycholate) with DNaseI for 15 minutes at room temperature followed by incubation with 2mL of the extraction buffer without DNaseI for an additional hour with gentle shaking. The extracted parasites were blocked with 5% BSA, 0.1% fish gelatin / PBS for 10 minutes at room temperature, rinsed once with incubation buffer (0.8% BSA, 0.1% fish gelatin in PBS), then incubated overnight with rabbit anti-TgDCX. Grids were then washed with PBS (3×5 min then 1×15 min), incubated with secondary antibody (anti-rabbit IgG conjugated with 1.4nm gold, Nanoprobes, Yaphank, New York, United States), diluted 1:160 in incubation buffer; incubated for 3 hours at room temperature then washed with PBS (3×5 min then 1×15 min). The sample was fixed for 5min with 1% glutaraldehyde in PBS and washed in distilled water 3 × 5 minutes. Silver enhancement was carried out using the HQ silver enhancement kit (Nanoprobes) by floating grids on a mixture of the initiator, activator and modulator for 2 minutes. The grid was washed with distilled water then stained using 2% phosphotungstic acid (pH7.0).

### Southern blot

Nick translation was used to synthesize single-stranded biotin-labeled probe DNA. The labeling reaction included 5 units of Klenow fragment (3’-5’ exo minus; NEB M0212), 3 µM biotinylated random octamers, 25 mM Tris-HCl, 100 mM HEPES, 1 mM DTT, 2.5 mM MgCl_2_, 0.1 mM dCTP, 0.1 mM dGTP, 0.1 mM dTTP, 0.084 mM dATP, 0.016 mM biotin-14-dATP, pH 7.0 @ 25°C. Template DNA (100–500ng) was denatured in boiling water for 5 min, put on ice for a few minutes, then incubated with the labeling reaction mix at 37°C for 6 hours. Labeled probe was isolated and purified by ethanol precipitation. Template DNA for the “exon 2–5” probe was made by PCR amplification of a 641 bp fragment of the TgDCX CDS, spanning the region from the 4th codon of exon 2 (M16) to the 2nd codon of exon VI (E225) using primers S5 and AS20. Template DNA for the 5’-UTR probe was a synthesized sequence corresponding to the 893 bp immediately preceding the first methionine codon of TgDCX in *T.gondii* genomic DNA.

~5 µg of *T. gondii* genomic DNA was digested with appropriate restriction enzymes (see Figure 7), run on an agarose gel with a biotinylated DNA ladder, transferred and UV cross-linked to positively charged nylon membrane. The blot was pre-hybridized for 1–2 hours, then hybridized for 12–18 hours with 10 ng/mL of biotinylated probe. Detection of bound probe utilized 1 µg/mL streptavidin (NEB N7021) and 0.1 µg/ml biotinylated alkaline phosphatase (Vector Labs B-2005-1). The chemiluminescent signal generated with substrate CDP-Star^®^ reagent (Roche 12-041-677-001) was recorded on Kodak blue sensitive X-ray film.

### Plaque assay and overall doubling time

Plaque assays were performed as previously described with some modifications (Liu et al., 2016). A total of 50, 100 or 200 parasites were added to each well of a 6-well plate containing a confluent HFF monolayer, and incubated at 37°C for 10 days undisturbed. Infected monolayers were then fixed with cold methanol for 10 min with gentle shaking, washed with DPBS, stained with 0.5% crystal violet in 20% methanol for 30 minutes, gently rinsed with ddH2O, air dried, and scanned.

An approximate value for the overall doubling time of parasite lines in culture was estimated from the ratio of inoculum volume to total volume of medium in the flask and the time required for complete lysis of the host cell monolayer. For example, inoculating 70 µL of culture supernatant from a freshly lysed flask of wild-type parasites into a new flask containing 4.0 mL of culture medium results in complete lysis of the new flask 44 ± 4 hours later. This is an amplification of 58 fold (= 4.07/0.07), i.e., 2^5.86^-fold, in 44 hours, corresponding to a doubling time of approximately 7.5 (= 44/5.86) hours. Note that the doubling time measured in this way is a composite of all the events of the lytic cycle as they occur in cultures, including time to invade, cell cycle time, time required for host cell lysis, and time spent searching for a new host cell.

### Host cell invasion assay

To maximize the proportion of vigorously active parasites, parasites were harvested when the host cell monolayer was less than 50% lysed. Initial inoculation of the cultures was timed so that all five parasite lines to be examined would be ready at the same time. The monolayers were scraped off the flask, suspended in culture medium, and passed through a 23G needle. The concentration of parasites was determined by counting in a hemocytometer, and 10^7^ parasites of each line were inoculated into a nearly confluent layer of African green monkey kidney epithelial cells (BS-C-1, ATCC CCL-26) growing in 35 mm plastic culture dishes with a glass coverslip bottom (MatTek). The dishes were incubated at 37 °C for 1 hour, washed twice with DPBS, then fixed with 3.7% formaldehyde, 0.06% glutaraldehyde, for 15 minutes. The dishes were then washed 5 times with DPBS and blocked with 1% BSA in DPBS for 30–45 minutes. Parasites that had attached to the host cell surface but had not yet invaded were labeled by incubation with a mouse antibody against SAG1 for 30–45 min, washed in DPBS 5 times, then incubated with Alexa568 goat anti-mouse. After permeabilization with 0.5% (v/v) Triton-X100 in DPBS for 15 min, washing 5 times with DPBS, and blocking for 30 min with 1% BSA in DPBS, all parasites were labeled by incubating with a rabbit antibody against SAG1 and a Cy2 goat anti-rabbit.

3D stacks of images of the red and green fluorescent parasites were acquired with a 20× NA 0.75 objective lens along with a reference DIC image from the middle of the stack. Images were collected at 15 different locations in each dish for each line, and the entire experiment was repeated three times. The fluorescent images were projected along the optic axis and deconvolved with a 2D optical transfer function for the objective lens. The image processing program Fiji (Schindelin et al., 2012; Schneider et al., 2012) was used to automate counting the red and green fluorescent parasites in each of the 450 images.

### Light microscopy

Parasites were grown on a subconfluent layer of HFF, rat aorta cells (A7r5, putative smooth muscle, ATCC CRL-1444) or BS-C-1 cells in a 35-mm plastic dish with #1.5 glass coverslip bottom (MatTek P35G-1.5–10-C). Just before transferring to the microscope, the medium was replaced with CO2-independent medium (Gibco 18045–088). The dish was maintained at 37°C in a humidified environmental chamber surrounding the microscope stage. 3D image stacks were collected at z-increments of 0.3 µm on an Applied Precision DeltaVision workstation using a 60× NA 1.3 silicone immersion lens. Brightly fluorescent 0.2 µm beads placed in the dish were used to adjust the correction collar of the lens to minimize spherical aberration. Deconvolved images were computed using the point-spread functions and software supplied by the manufacturer. SIM images were collected at z-increments of 0.125 µm on an Applied Precision OMX 3D-SIM system using a 100X NA 1.4 objective lens, and processed using the manufacturer’s software. All quantitative measurements were carried out using the raw image data, but in the final displayed images, the relative brightness of different fluorophores has been altered as needed to best utilize the gamut of a computer monitor LCD display.

To image the fluorescent virus particles, 7 µL of virus culture supernatant or sucrose-gradient purified virus was spread on a cleaned 22×22 mm #1.5 coverslip, allowed to absorb for 2 min, then inverted on a 3 µL droplet of buffer (50 mM Na-HEPES pH 7.0, 0.1 M NaCl, 0.5 mM Na_2_EDTA) on a cleaned glass slide. Excess liquid was removed by pressing the slide, coverslip down, onto filter paper for a few seconds. The edges of the coverslip were sealed with VALAP (Vaseline:Lanolin:Paraffin wax 1:1:1) or nail polish. Images were acquired on an Applied Precision DeltaVision workstation using a 60x NA 1.3 silicone immersion lens.

### Quantitative intensity comparisons of Sindbis virions and Toxoplasma organelles

All image analysis for quantitative intensity comparisons was carried out using an updated version of the Semper (Saxton et al., 1979) software package (source code generously provided by Dr. Owen Saxton) running under Unix on a Mac Mini or MacBookPro. Pre-processing steps included defining the portion of the field of view over which the illumination intensity varied by less than 15%, and then correcting for the non-uniform illumination across this restricted field of view (“flat-fielding”). Normalization of all images to a global average illumination intensity was performed, based on photosensor measurement of actual illumination intensity during each individual image acquisition (hardware and software built in to the Applied Precision Deltavision system). To help define the plane of best focus, 3D-stacks were deconvolved by constrained iterative deconvolution, using software supplied with the Applied Precision Deltavision microscope system. However, un-deconvolved single optical sections corresponding to the plane of best focus were used for analysis. Prior to quantitative analysis, images intensities in grey-levels were converted to intensities in photons/sec, using the known exposure time and an experimentally determined ADU gain calibration (Faruqi et al., 1999; Murray, 2007).

The images to be considered are characterized by well-defined small peaks of relatively high intensity surrounded by large areas of much lower intensity. Thus for each image, a local background can be defined by first excluding all areas with intensity exceeding a certain threshold, usually taken as five times the expected standard deviation of the average background assuming Poisson statistics. In images of fluorescent virus, which typically included >500 nearly identical particles per field of view, virus particles were identified automatically, making use of the extensive particle analysis tools within Semper. For images of organelles within *Toxoplasma*, which generally included <20 organelles with more variable intensity per field of view, organelles were first identified manually on the display using a cursor, and subsequently processed automatically as for the virus particles. Regions of interest surrounding each identified virus particle or organelle were defined as contiguous sets of pixels with intensity exceeding the local background by some threshold, typically ten times the expected Poisson standard deviation of the local background. Pixels within each (possibly 3D) region of interest so defined were then counted, their intensities summed, and the local background intensity multiplied by the number of pixels within the ROI was subtracted to give the net integrated intensity for each virus or organelle.

### Electron microscopy of whole mount Toxoplasma

Parasites from 1 mL of a freshly lysed culture were collected by centrifugation for 1 min at 3600 × g, washed once with HEPES-buffered serum-free culture medium, and resuspended in 40 µL of the same medium containing 20 µM calcium ionophore A23187. After 2 min, 4 µL of parasite suspension was pipetted onto a carbon-coated EM grid and placed in a humid chamber for 8 minutes. The grid was then placed parasite-side down onto a 50 µL drop of 0.5% Triton-X100 for 3 min, transferred to a drop of 0.002% Triton-X100, 2% phosphotungstic acid pH 7.3 for 2 min, and all of the liquid was wicked off by touching the grid to the edge of a piece of filter paper. Images were collected with a JEOL 1010 electron microscope operated at 80 kev.

## ACKNOWLEDGEMENTS

We thank Dr. Lloyd Kapser (Dartmouth College, Hanover, NH) for the mouse anti-SAG1 antibody, Dr. Richard Day (Indiana University School of Medicine, Indianapolis, IN) for a plasmid containing the mNeonGreenFP coding sequence, Dr. Suchetana Mukhopadhyay (Indiana University, Bloomington IN) for a plasmid encoding the complete genome of recombinant Sindbis virus TE12, Dr. Jacqueline Leung (Indiana University) for insightful discussions, and Amanda Rollins for technical support. This study was supported by funding from the March of Dimes (6-FY15-198), NIH-NIAID (R01-AI098686) awarded to KH, and facility funding from the Indiana Clinical and Translational Sciences Institute to KH, funded in part by grants #UL1 TR001108 and #TL1TR001107 from the National Institutes of Health, National Center for Advancing Translational Sciences, Clinical and Translational Sciences Award.

## Bibliography

Amos, L. A. and Amos, W. B. (1991). The bending of sliding microtubules imaged by confocal light microscopy and negative stain electron microscopy. J.Cell.Sci.Suppl. 14, 95–101.

Carey, K. L., Westwood, N. J., Mitchison, T. J. and Ward, G. E. (2004). A small-molecule approach to studying invasive mechanisms of Toxoplasma gondii. Proc Natl Acad Sci U S A 101, 7433–8.

de Leon, J. C., Scheumann, N., Beatty, W., Beck, J. R., Tran, J. Q., Yau, C., Bradley, P. J., Gull, K., Wickstead, B. and Morrissette, N. S. (2013). A SAS-6-like protein suggests that the Toxoplasma conoid complex evolved from flagellar components. Eukaryot Cell 12, 1009–19.

Desser, S. S. (1970). The fine structure of Leucocytozoon simondi. III. The ookinete and mature sporozoite. Canadian Journal of Zoology 48, 641–645.

Faruqi, A. R., Henderson, R. and Subramaniam, S. (1999). Cooled CCD detector with tapered fibre optics for recording electron diffraction patterns. Ultramicroscopy 75, 235–250.

Gleeson, J. G., Lin, P. T., Flanagan, L. A. and Walsh, C. A. (1999). Doublecortin is a microtubule-associated protein and is expressed widely by migrating neurons. Neuron 23, 257–71.

Heaslip, A. T., Ems-McClung, S. C. and Hu, K. (2009). TgICMAP1 is a novel microtubule binding protein in Toxoplasma gondii. PLoS ONE 4, e7406.

Heaslip, A. T., Dzierszinski, F., Stein, B. and Hu, K. (2010). TgMORN1 is a key organizer for the basal complex of Toxoplasma gondii. PLoS Pathog 6, e1000754.

Heaslip, A. T., Nishi, M., Stein, B. and Hu, K. (2011). The motility of a human parasite, Toxoplasma gondii, is regulated by a novel lysine methyltransferase. PLoS Pathog 7, e1002201.

Hernandez, R., Sinodis, C. and Brown, D. T. (2010). Sindbis virus: propagation, quantification, and storage. Curr Protoc Microbiol **Chapter 15**, Unit15B 1.

Hlavanda, E., Kovacs, J., Olah, J., Orosz, F., Medzihradszky, K. F. and Ovadi, J. (2002). Brain-specific p25 protein binds to tubulin and microtubules and induces aberrant microtubule assemblies at substoichiometric concentrations. Biochemistry 41, 8657–64.

Hu, K. (2002). Building a parasite: The study of the cell division and cytoskeleton of *Toxoplasma gondii*. In Biology, pp. 187. Philadelphia: University of Pennsylvania.

Hu, K., Roos, D. S. and Murray, J. M. (2002). A novel polymer of tubulin forms the conoid of Toxoplasma gondii. Journal of Cell Biology 156, 1039–50.

Hu, K., Suravajjala, S., DiLullo, C., Roos, D. S. and Murray, J. M. (2003). Identification and characterization of new tubulin isoforms in *Toxoplasma gondii*. In American Society for Cell Biology 43rd Annual Meeting. San Francisco, CA.

Hu, K., Roos, D. S., Angel, S. O. and Murray, J. M. (2004). Variability and heritability of cell division pathways in Toxoplasma gondii. Journal of Cell Science 117, 5697–705.

Hu, K., Johnson, J., Florens, L., Fraunholz, M., Suravajjala, S., DiLullo, C., Yates, J., Roos, D. S. and Murray, J. M. (2006). Cytoskeletal components of an invasion machine--the apical complex of Toxoplasma gondii. PLoS Pathog 2, 121–138.

Hu, K. (2008). Organizational changes of the daughter basal complex during the parasite replication of Toxoplasma gondii. PLoS Pathog 4, e10.

Jose, J., Tang, J., Taylor, A. B., Baker, T. S. and Kuhn, R. J. (2015). Fluorescent Protein-Tagged Sindbis Virus E2 Glycoprotein Allows Single Particle Analysis of Virus Budding from Live Cells. Viruses 7, 6182–99.

Kim, M. H., Cierpicki, T., Derewenda, U., Krowarsch, D., Feng, Y., Devedjiev, Y., Dauter, Z., Walsh, C. A., Otlewski, J., Bushweller, J. H. et al. (2003). The DCX-domain tandems of doublecortin and doublecortin-like kinase. Nature Structural Biology 10, 324–33.

Kurachi, M., Hoshi, M. and Tashiro, H. (1995). Buckling of a single microtubule by optical trapping forces: direct measurement of microtubule rigidity. Cell Motility and the Cytoskeleton 30, 221–8.

Leander, B. S. and Keeling, P. J. (2003). Morphostasis in alveolate evolution. Trends in Ecology & Evolution 18, 395–402.

Leveque, M. F., Berry, L. and Besteiro, S. (2016). An evolutionarily conserved SSNA1/DIP13 homologue is a component of both basal and apical complexes of Toxoplasma gondii. Scientific Reports 6, 27809.

Liliom, K., Wagner, G., Kovacs, J., Comin, B., Cascante, M., Orosz, F. and Ovadi, J. (1999). Combined enhancement of microtubule assembly and glucose metabolism in neuronal systems in vitro: decreased sensitivity to copper toxicity. Biochemical and Biophysical Research Communications 264, 605–10.

Liu, J., He, Y., Benmerzouga, I., Sullivan, W. J., Jr., Morrissette, N. S., Murray, J. M. and Hu, K. (2016). An ensemble of specifically targeted proteins stabilizes cortical microtubules in the human parasite Toxoplasma gondii. Molecular Biology of the Cell 27, 549–71.

Lustig, S., Jackson, A. C., Hahn, C. S., Griffin, D. E., Strauss, E. G. and Strauss, J. H. (1988). Molecular basis of Sindbis virus neurovirulence in mice. Journal of Virology 62, 2329–36.

Moores, C. A., Perderiset, M., Francis, F., Chelly, J., Houdusse, A. and Milligan, R. A. (2004). Mechanism of microtubule stabilization by doublecortin. Mol Cell 14, 833–9.

Murray, J. M. (2007). Practical aspects of quantitative confocal microscopy. Methods Cell Biol 81, 467–78.

Nagayasu, E., Zhang, F., Hu, K., Ananvoranich, S. and Murray, J. M. (2006). Cytoskeletal components of an invasion machine: The apical complex and conoid of *Toxoplasma gondii*. In American Society for Cell Biology 46th Annual Meeting, vol. 22, pp. 293. San Diego, CA.

Nishi, M., Hu, K., Murray, J. M. and Roos, D. S. (2008). Organellar dynamics during the cell cycle of Toxoplasma gondii. Journal of Cell Science 121, 1559–68.

Orosz, F. (2009). Apicortin, a unique protein, with a putative cytoskeletal role, shared only by apicomplexan parasites and the placozoan Trichoplax adhaerens. Infect Genet Evol.

Orosz, F. (2016). Wider than Thought Phylogenetic Occurrence of Apicortin, A Characteristic Protein of Apicomplexan Parasites. Journal of Molecular Evolution 82, 303–314.

Patra, K. P. and Vinetz, J. M. (2012). New ultrastructural analysis of the invasive apparatus of the Plasmodium ookinete. American Journal of Tropical Medicine and Hygiene 87, 412–7.

Portman, N., Foster, C., Walker, G. and Slapeta, J. (2014). Evidence of intraflagellar transport and apical complex formation in a free-living relative of the apicomplexa. Eukaryot Cell 13, 10–20.

Portman, N. and Slapeta, J. (2014). The flagellar contribution to the apical complex: a new tool for the eukaryotic Swiss Army knife? Trends Parasitol 30, 58–64.

Rice, C. M., Levis, R., Strauss, J. H. and Huang, H. V. (1987). Production of infectious RNA transcripts from Sindbis virus cDNA clones: mapping of lethal mutations, rescue of a temperature-sensitive marker, and in vitro mutagenesis to generate defined mutants. Journal of Virology 61, 3809–19.

Roos, D. S., Donald, R. G., Morrissette, N. S. and Moulton, A. L. (1994). Molecular tools for genetic dissection of the protozoan parasite Toxoplasma gondii. Methods in Cell Biology 45, 27–63.

Schindelin, J., Arganda-Carreras, I., Frise, E., Kaynig, V., Longair, M., Pietzsch, T., Preibisch, S., Rueden, C., Saalfeld, S., Schmid, B. et al. (2012). Fiji: an open-source platform for biological-image analysis. Nat Methods 9, 676–82.

Schneider, C. A., Rasband, W. S. and Eliceiri, K. W. (2012). NIH Image to ImageJ: 25 years of image analysis. Nat Methods 9, 671–5.

Shaner, N. C., Lambert, G. G., Chammas, A., Ni, Y., Cranfill, P. J., Baird, M. A., Sell, B. R., Allen, J. R., Day, R. N., Israelsson, M. et al. (2013). A bright monomeric green fluorescent protein derived from Branchiostoma lanceolatum. Nat Methods 10, 407–9.

Verdaasdonk, J. S., Lawrimore, J. and Bloom, K. (2014). Determining absolute protein numbers by quantitative fluorescence microscopy. Methods in Cell Biology 123, 347–65.

